# Time-course of population activity along the dorsoventral extent of the superior colliculus during delayed saccade tasks

**DOI:** 10.1101/571307

**Authors:** Corentin Massot, Uday K. Jagadisan, Neeraj J. Gandhi

## Abstract

The superior colliculus (SC) is an excellent substrate to study functional organization of sensorimotor transformations. We used linear multi-contact array recordings to analyze the spatial and temporal properties of population activity along the SC dorsoventral axis during delayed saccade tasks. During the visual epoch, information appeared first in dorsal layers and systematically later in ventral layers. In the ensuing delay period, the laminar organization of low-spiking rate activity matched that of the visual epoch. During the pre-saccadic epoch, spiking activity emerged first in a more ventral layer, ∼100ms before saccade onset. This buildup of activity appeared later on nearby neurons situated both dorsally and ventrally, culminating in a synchronous burst across the dorsoventral axis, ∼28ms before saccade onset. Stimulation of individual contacts on the laminar probe produced saccades of similar vectors. Collectively, the results reveal a principled spatiotemporal organization of SC population activity underlying sensorimotor transformation for the control of gaze.

## Introduction

Our interactions with the environment are mediated via brain networks that transform sensory signals to motor actions at the appropriate time. In the context of gaze control, this sensorimotor transformation entails processing of incoming visual information and generating a movement command to appropriately redirect the line of sight. The superior colliculus (SC) in the midbrain modulates its activity in response to both stimulus presentation and movement generation, as well as during the interval between the two events. Like cortex, the SC is composed of distinct layers. Its superficial layers are predominantly driven by visual processing structures like the retina and primary visual cortex, while its deeper layers communicate a broad spectrum of information with many cortical and noncortical areas ^1-3^. It also has a canonical organization with established microcircuits for communication both within and across layers ^4^. Finally, it has a topographic representation of visual space and for generation of gaze shifts to those locations ^5^. Thus, the SC is ideally suited to study the neural correlates of sensorimotor transformation.

Despite a wealth of knowledge about the anatomical organization of the SC and the functional properties of individual SC neurons, current understanding about the link between structural and functional organization at the population level is limited. For instance, it is unclear whether the properties of individual neurons exhibit systematic spatiotemporal organization during the sensorimotor transformation, and how such organization is linked to the microarchitecture of the SC network. Bridging structure with function helps to not only understand the computations underlying the transformation but also to build biologically-inspired network models of sensorimotor learning and behavior ^6,7^.

Linear microelectrodes have recently been used for such structure-function mapping, particularly in regions on the cortical surface, since they enable the simultaneous measurement of neural activity across multiple layers. Indeed, this approach has provided insights into how sensory ^8-11^ and cognitive processes like spatial attention ^12, 13^, working memory ^14^, decision-making ^15^, and episodic encoding ^16^ are mediated as a function of depth as well as about modes of communication between layers in driven and quiescent states 17,18. In our case, sensorimotor transformations occurring within a single brain region provide a unique opportunity to study the link between structural organization, functional physiology, and behavior. Thus, we extended the use of the laminar probe to the SC in the subcortex to investigate the visual to motor transformation as a function of depth. Our electrode penetrations were approximately orthogonal to SC surface and hence encountered neurons that responded vigorously for the same sensory and motor vectors. We were therefore able to test whether SC neurons exhibit fine-grained spatial and temporal organization that is particularly suited to implement the sensorimotor transformation

We recorded from the SC of two monkeys performing delayed saccade tasks with linear, multi-contact probes. We used current source density (CSD) analyses to obtain a veridical estimate of the relative probe depth in SC during any given penetration and align data from multiple sessions ^13,18^. We found a strong and systematic depth-dependent organization for both intensity and timing of neural activity. Neurons across layers exhibited both visual and movement-related responses, but visual-preferring neurons were more likely to reside in dorsal layers, with a gradual, non-linear transformation to movement-preference occurring at deeper sites. The majority of SC neurons modulated their firing rates during both visual and movement epochs. The latency of the visual responses increased monotonically with depth. The activity during the delay period decreased to a low-spiking rate but was still higher for dorsal layers. In contrast, pre-saccadic buildup activity originated at intermediate depths and systematically spread bidirectionally in dorsal and ventral neurons. Buildup activity culminated in a punctate saccade-related burst that was synchronized across all neurons along the dorsoventral axis. These results reveal important spatiotemporal patterns of activity organization that advance our understanding of the neural network activity within SC. We present these results in the context of other studies of functional organization and discuss the potential implications of a structure-to-function mapping for sensorimotor transformations.

## Results

Multiunit spiking activity (MUA) and local field potentials (LFPs) were recorded on each contact of a 16-channel laminar probe that spanned the dorsoventral extent of the SC in two Rhesus monkeys performing visually-guided (VG) and memory-guided (MG) delayed saccade tasks (Figure 1). A total of 26 sessions were recorded from 16 different grid locations in the chamber. Of these, 20 were included for the analysis of the VG task. 3 sessions were excluded because of noisy LFP signals and 3, because of poor signal-to-noise ratio of spiking activity. 13 of the 26 sessions were also recorded with the MG task. 2 of these sessions were excluded because of poor signal-to-noise ratio. The intended angle of penetration was orthogonal to the SC surface so that all electrode contacts encountered neurons with similar preferred visual and/or motor vectors.

**Figure 1.**
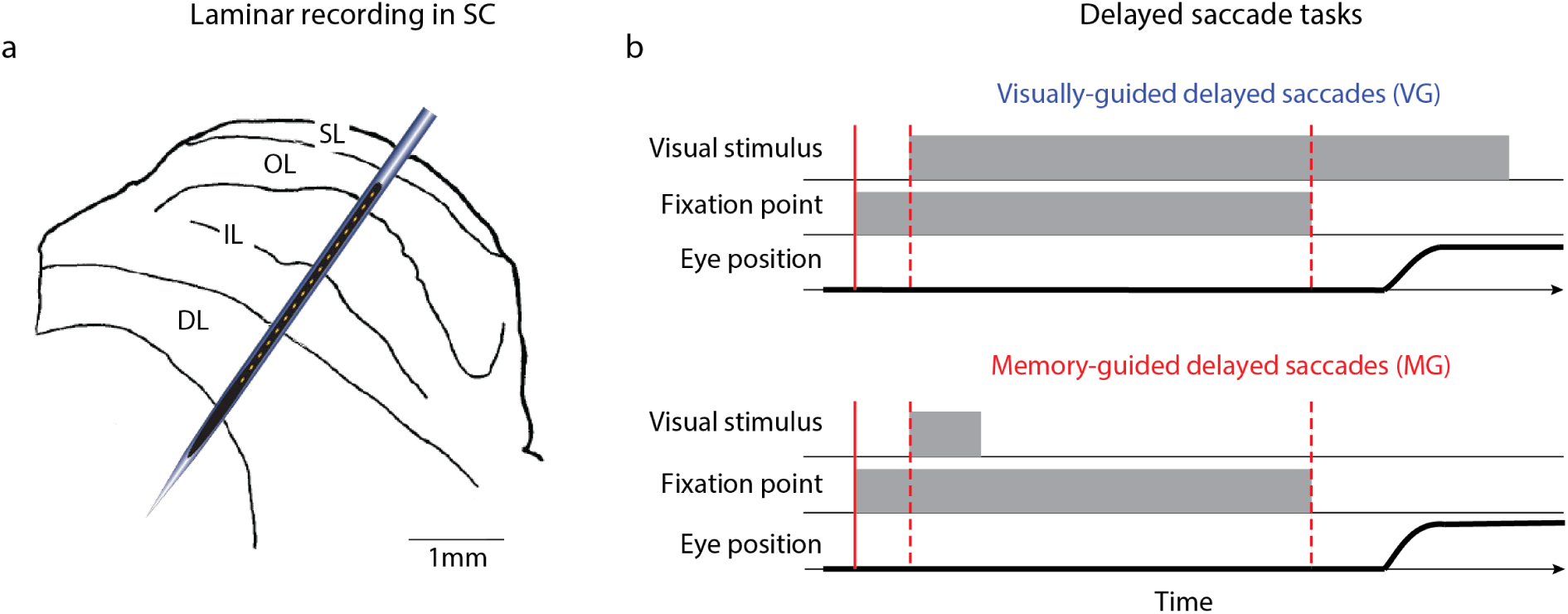
Laminar recording across the dorsoventral extent of SC. (a) Schematic of SC laminar structure and laminar probe. The probe (150µm inter-contact distance; ∼300µm diameter) is drawn roughly to scale with the SC sketch. SL (superficial layers), OL (optic layers), IL (intermediate layers), DL (deep layers) (modified from ^79^). (b) Visual illustrations of the visually-guided (VG) and memory-guided (MG) delayed saccade tasks. See text for details.

### Laminar organization of activity levels

Spike density functions aligned on visual burst and saccade onsets for an individual session of VG trials are shown in Figure 2a-b. Each trace is an average across trials for which the stimulus location matched the optimal vector estimated for the penetration (see *Materials and Methods*). The waveform on each contact is scaled and shifted vertically for visualization. All channels increased activity in response to the visual target and in most cases showed two peaks. All movement bursts started before and peaked around saccade onset. Some movement bursts also displayed a second peak ∼50ms after saccade onset, which was attributed to a post-saccadic visual response. These are well-known characteristics of SC neural activity ^19-21^.

**Figure 2.**
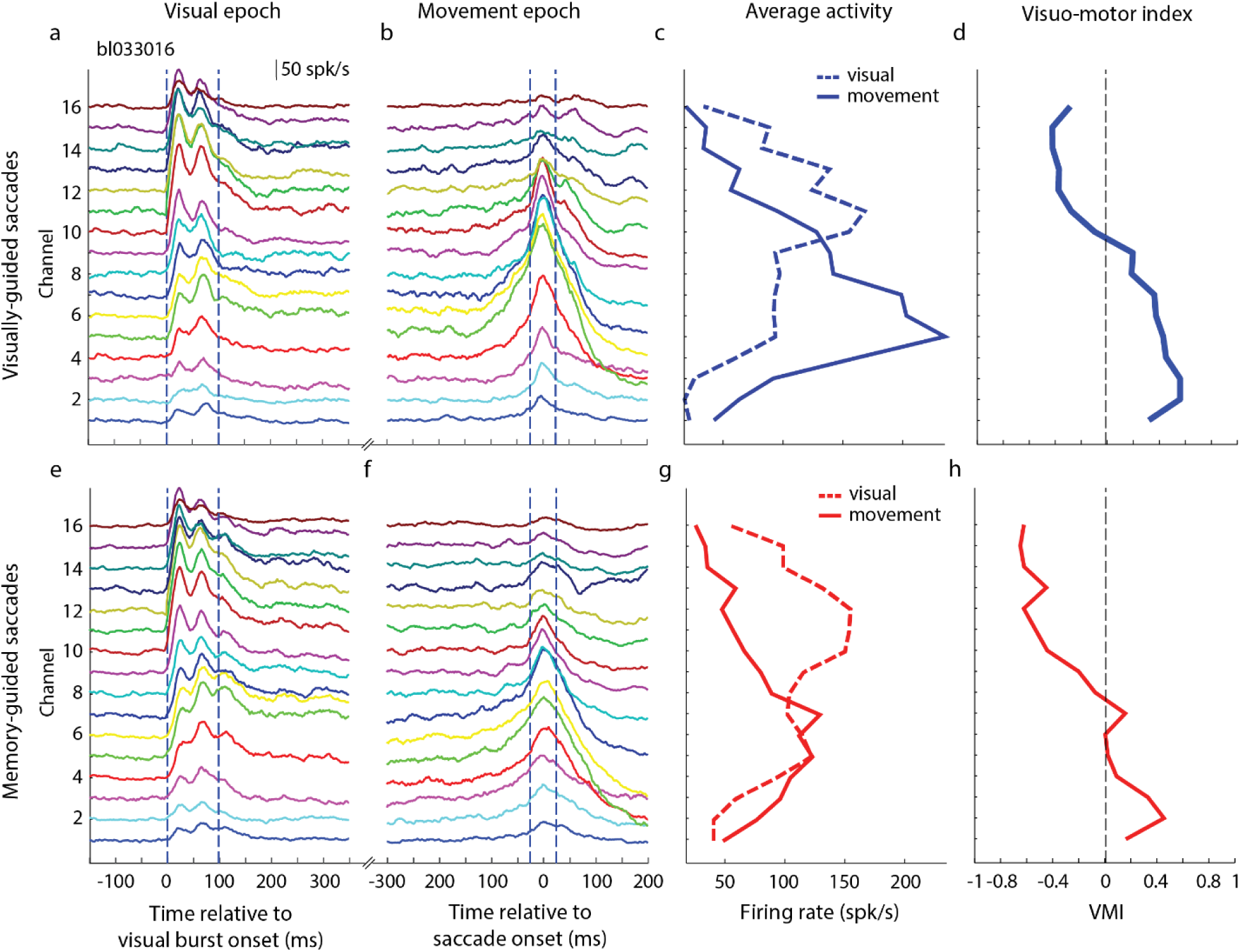
Example data of laminar recording from a single session. (a) Plots of trial-averaged spike density waveforms for each channel, aligned on visual burst onset (*t* = 0 *ms*). The method for aligning on burst is described in Methods section. The waveform on each contact is scaled and shifted vertically for visualization. The dashed vertical blue lines show the boundaries of the visual epochs used in panel c. Data are from VG task. (b) Trial-averaged waveforms aligned on saccade onset, using the same convention used in panel a. (c) The average firing rates during the visual (dashed blue line) and movement (solid blue line) epochs are plotted as a function of depth. (d) The visuomotor index (VMI) is plotted as a function of depth. Negative and positive VMI values denote greater visual and movement related activities, respectively. Vertical line denotes VMI=0. (e-h) Data for MG trials from the same session. Display convention is same as for panels (a-d).

A previously under-appreciated feature that emerged from the laminar probe approach is the systematic pattern of activity levels dorsoventrally along the penetration. The average firing rate in the visual epoch increased gradually with depth, reached a maximum at a relatively dorsal site (contact 11 for this example, Figure 2c), and then decreased again for more ventral sites. The activity during the movement epoch followed the same trend except that it was shifted and peaked at a more ventral site (contact 5; contact 1 being the most ventral). We quantified the contrast between the visual and the movement related bursts by computing a visuo-movement index (VMI) (see *Materials and Methods*). The index was negative (dominated by the visual response) for the most dorsal channels and gradually became positive (dominated by the movement response) for increasingly more ventral channels, before plateauing for the most ventral channels (Figure 2d). Figure 2e-h reports the same results for MG trials from the same session. The trends in spike density waveforms across depths, the average firing rates in the visual and movement epochs, and the VMI are very similar to VG trials. One notable difference, as reported previously, is that the movement burst is attenuated in MG trials ^22^, particularly for the more ventral channels. This is also reflected by a slight diminution of the VMI relative to VG trials for the ventral channels.

We next sought to average the similar trends we observed across sessions. To do so, we first needed a method to properly align the data to a reference contact or depth. One consideration is to use microdrive readings, which corresponds to the absolute depth of the linear probe from the dura. However, it is not a reliable measure because of two main limitations. First the daily setup of the recording equipment introduces slight configuration changes that are difficult to control for (notably the initial position of the probe). Second the viscosity of the cerebral tissue, which can change both within and across sessions, introduces an inherent variability. This makes it unlikely that the same absolute depth of the probe, as indicated by the micro-drive, corresponds to the same relative position within SC. While the effects of the recording setup can be mitigated ^23^, the viscosity of the tissue cannot be controlled for. To overcome these limitations, we used an objective method based on features in the current-source density analysis (CSD) of the LFP signals from the visual epoch. Supplementary Figure 1 presents the CSD analysis and the different steps of the alignment procedure (see *Materials and Methods*).

The outcome of CSD-guided alignment and averaging across sessions for firing rates in the visual and movement epochs are shown in Figure 3. For VG trials (panel a), the amplitude of the visual activity plateaus from channel 2 to 6 peaks with a peak at channel 4 at 88.2spk/s (95%CI [56.1 130.3]spk/s), and gradually decreases for the other dorsal and ventral channels. The amplitude of the movement activity peaks at channel −2 at 136.7spk/s (95%CI [112.0 162.3]spk/s), and gradually decreases in both dorsal and ventral directions. These general trends are similar for MG trials (panel b). The amplitude of the visual activity plateaus from channel 2 to 6 with a peak at channel 6 at 92.3spk/s (95%CI [57.8 139.2]spk/s) and gradually decreases for the other dorsal and ventral channels. The amplitude of the movement activity peaks at channel −1 at 113.7spk/s (95%CI [86.5 140.4]spk/s), and gradually decreases for the other dorsal and ventral channels. The most notable difference between VG and MG trials is the smaller, but not significantly different (Wilcoxon rank-sum test, P>0.067, for all channels), peak amplitude of movement activity for MG trials. Overall these results show that the amplitudes of two bursts are systematically organized across depths. The peak visual activity is situated between 0.75mm and 1.05mm (between channels 5 and 7) more dorsally than the peak movement activity, reflecting a visual preference for dorsal channels and movement preference for ventral channels.

**Figure 3.**
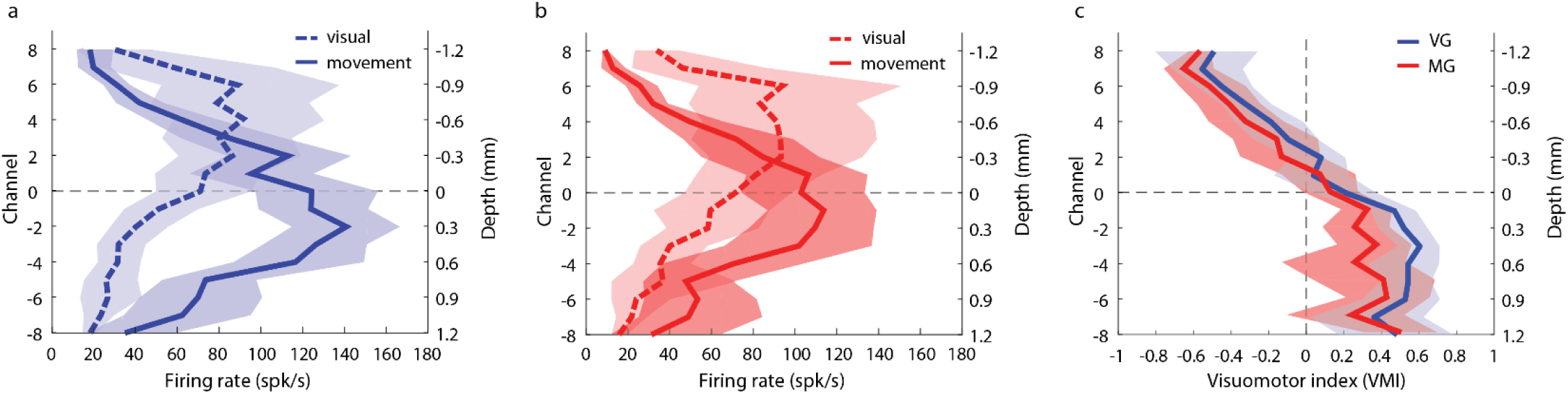
Population-averaged visual and motor activity. The peak firing rate averaged across sessions during the visual (dashed trace) and movement (solid traces) are plotted as a function of channel number or depth for VG (panel a; blue traces) and MG (panel b; red traces) tasks. (c) The session-averaged VMI is plotted against depth for VG (solid blue line) and MG (solid red line) trials. For all panels, the channel index is reported on the left ordinate axis. Channel 0 corresponds to the CSD reference channel that was used to align the data across sessions (see *Materials and Methods*). The right ordinate axis indicates the relative depth in mm. The blue and red translucid regions surrounding same color traces in all three panels represent 95% confidence interval of the VG and MG trials, respectively.

We next evaluated whether the relative contributions of the visual and movement activity on each channel also follow a systematic organization along the dorsoventral axis of the SC. We thus measured the contrast between the visual and the movement amplitude activity by computing a visuo-movement index (VMI) (see *Materials and Methods*). The VMI analysis across all sessions reinforces the trends observed in the two distributions (Figure 3c). The contrast ratio was linear from dorsal channels down to the reference channel, but then plateaued for deeper channels. It is important to note that the plateau was not due to constant firing rates of visual and movement bursts. In fact, both visual and movement activities decreased ventrally from the reference channel, however their relative amplitudes remained constant, with the movement activity being higher. The VMI trends were similar for both VG and MG trials, as evidenced by the superposition of the confidence interval (CI) bands during the linear part. Ventrally to the reference channel, the plateau values are slightly different (∼0.5, VG trials; ∼0.4, MG trials) but not statistically significantly (Wilcoxon rank sum test, P>0.033, for all channels). This is due to the lower firing rate of movement activity for MG trials ^22^.

In addition to quantitative parameters like VMI, SC neurons are also parsed into visual, visuomotor (or visuo-movement), and motor (or movement) classifications. We too categorized each neuron’s activity based on the significance of their visual and movement activity (see *Materials and Methods*). Of the whole population of recorded channels with significant MUA activity, visuo-movement neurons constituted the majority (Figure 4c). Figure 4a shows the distribution of neurons in each category as a function of depth for VG trials. The neuron count, plotted on the abscissa, is normalized to the number of sessions (left panel) and shows that more neurons were sampled in the ventral part of SC. The neuron count is also normalized individually for each channel to the number of significant MUA (right panel), in order to compensate for the difference in recorded MUA across channels. Visual-only MUA were mainly found at the most dorsal channels (channels 6, 7 and 8) and represented only a small proportion of the number of recorded units. This could be a consequence of difficulty isolating these smaller neurons ^1^. Visuo-movement MUA were found across all depths and was the dominant category between channels 0 and 6. Movement-only MUA were mainly found on channels ventral to the reference channel and their proportion increased with depth. Figure 4b shows the categorization of MUA for MG trials. The distributions were qualitatively similar to VG trials. These results collectively confirm the existing view, obtained from single electrode experiments, that the activity in SC is not randomly distributed across depths but instead follows a general principle: visual activity is predominant at dorsal depths and movement activity at ventral depths; in between, both visual and movement are mixed within the same MUA. Another crucial observation that emerges from this analysis is that SC neurons with the most vigorous movement-related burst reside in the visuo-movement, not movement, neuron category. This result has important consequence on how brainstem neurons that receive SC activity identify or decode the burst (visual or movement epoch) that triggers the movement (see *Discussion*).

**Figure 4.**
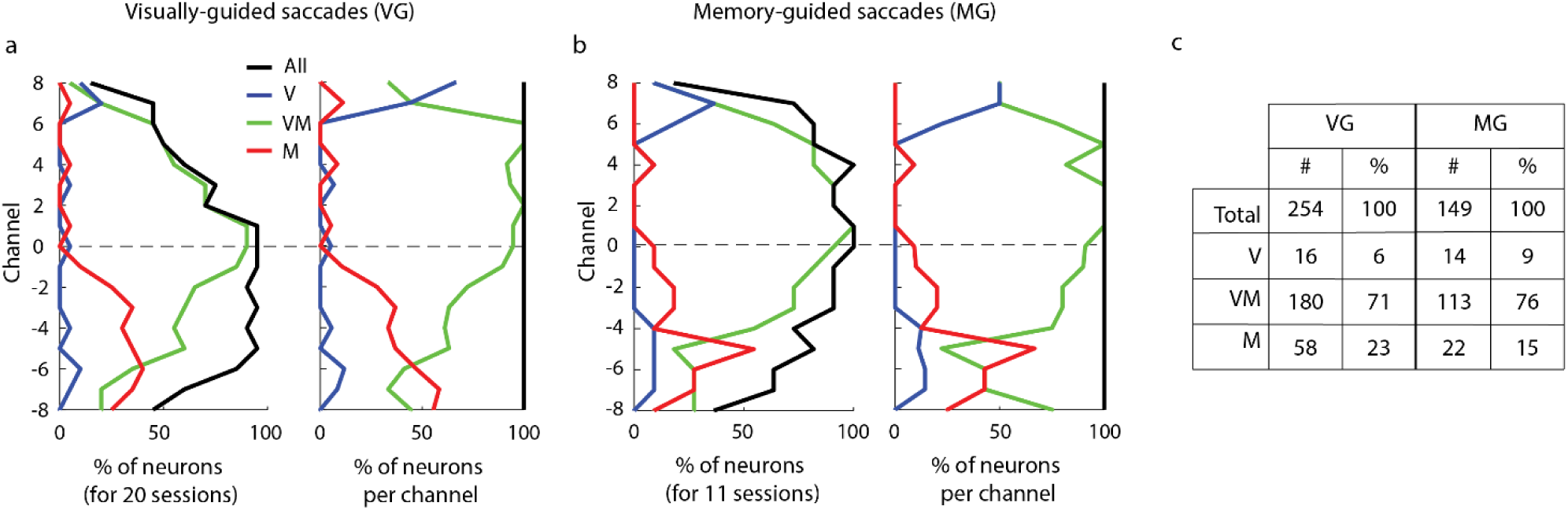
Categorization of SC neurons. Every panel shows the distributions of visual-only (V, blue trace), visuo-movement (VM, green trace) and movement-only (M, red trace) neurons as a function of channel number. Data are pooled across 20 VG sessions (a) and 11 MG sessions (b). In each subplot, the abscissa denotes the proportion of neurons. Left: For each channel, the neuron count (either for each category or for all neurons) across all sessions is normalized by the number of sessions (20 for VG and 11 for MG trials). Right: The neuron count is normalized individually for each channel to compensate for the non-uniform sampling of neurons across depths (i.e., the neuron count becomes 1 on every channel). (c) A summary of the percentages of each type of neuron for VG and MG trials.

We next evaluated the distribution of delay period activity as a function of depth for VG and MG trials (Figure 5). Each color represents the across-sessions average of the baseline-corrected delay activity observed in nonoverlapping 50ms time bins, starting from the last peak of the visual burst across channels (blue, ‘Bin 1’) to the end of the shortest delay period (orange, ‘Bin 6’). The activity was, as expected, highest in the wake of the visual burst and then decreased gradually to a low-spiking rate as the delay period progresses. The strongest response stayed consistently between channel 2 and 4 throughout the delay period for both types of trials. For VG trials, the activity reached 33.0spk/s (95%CI [14.5 61.1]spk/s) on channel 2 on bin 6. For MG trials, the activity reached 43.6spk/s (95%CI [13.0 83.8]spk/s) on channel 3 on bin 5. Notably, these channels were the same that discharged maximally for the visual burst (thick black trace). For the deepest channels, we noticed a difference between the two tasks. For MG tasks, the activity was slightly higher than the peak activity of the visual burst while it remained at minimum level for VG tasks. Note that the confidence intervals were very large and overlapping across bins which makes this increase of activity not statistically significant in our data. Further investigation is needed to reveal the possible cause and role of this increase for MG tasks.

**Figure 5.**
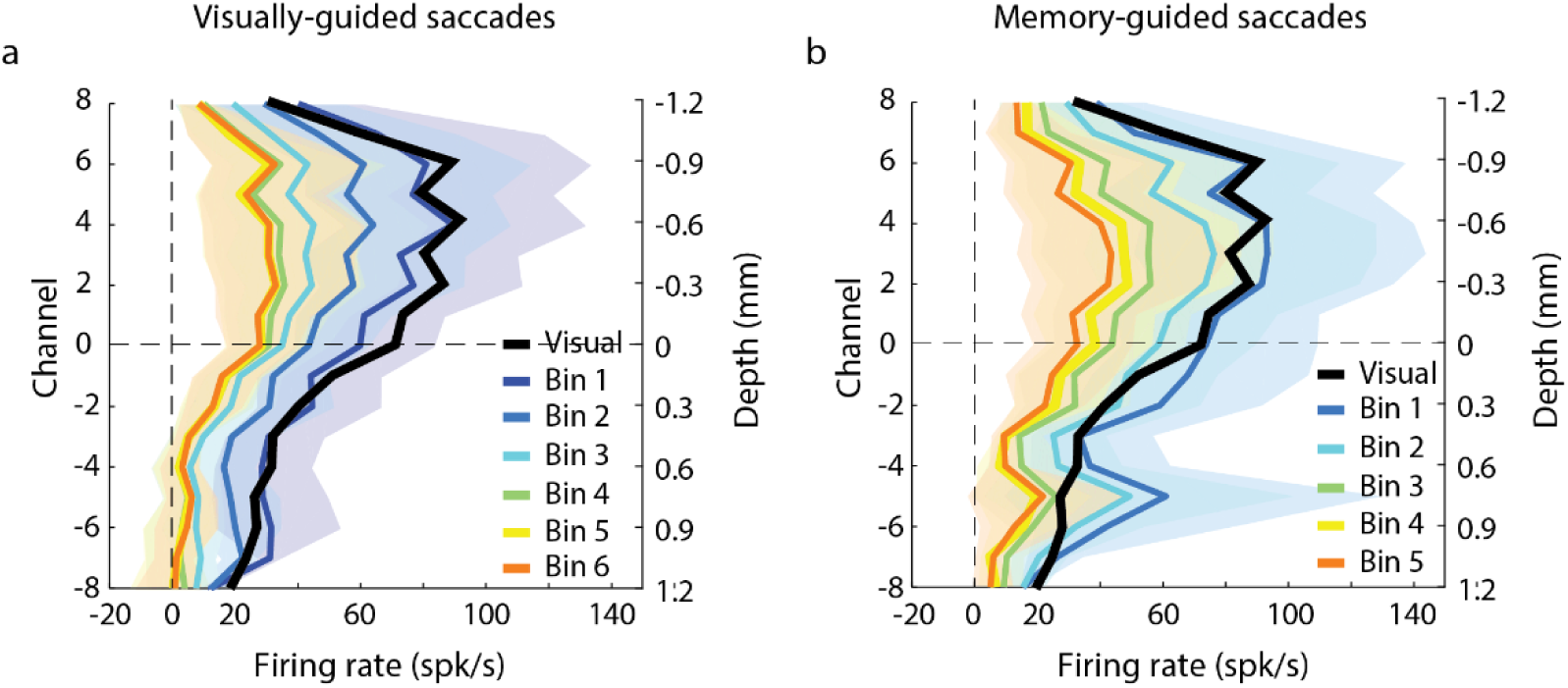
Population-averaged delay period activity within SC. Each trace represents mean activity in a specific interval as a function of depth and averaged across sessions for (a) VG and (b) MG tasks. The black traces is the mean activity in the visual epoch; it is identical to the dashed traces in Figure 3a,b. The remaining traces are averages computed over nonoverlapping 50ms bins starting from the last peak of the visual burst across channels (blue, ‘Bin 1’) to the end of the shortest delay period (orange, ‘Bin 6’). The colored translucid regions surrounding the thick traces represent the 95% confidence interval computed across sessions.

### Laminar organization of temporal events

We next examined the time-course of activity during the visual and the pre-saccadic epochs in order to identify potential spatiotemporal patterns along the dorsoventral axis. We first focused on the latency of the sensory response (visual burst) across depths. Data recorded during the visual epoch are typically aligned on target onset. With this type of alignment, the visual response appears 50-100ms after target onset. However, individual neurons and local circuitry response are stochastic. A delay in this range along with the associated variability could be large enough to average out latency differences across channels. To circumvent this concern, we aligned data on the onset of the visual burst. Briefly, for each trial and for each channel, burst onset was estimated using the *Poisson surprise* detection method ^24^. The channel with the most (not necessarily the earliest) detected bursts, henceforth referred to as the visual alignment channel (see *Materials and methods*), was used to align data on all channels. Figure 6a shows an example session (same one as Figure 2) with the spiking activity aligned on visual burst across; channel 11 was the visual alignment channel. Each colored, vertical mark indicates the onset of the visual activity, estimated using a 2-piecewise regression-based method (see Supplementary Figure 2 and *Materials and methods*). Figure 6b plots the visual latency estimate of each channel centered on the reference channel 7, i.e., the channel obtained from the CSD analysis for aligning data in depths across sessions. Note that the reference and alignment channels need not be the same. A cubic polynomial fit of the visual latencies reveals a general spatiotemporal trend for this session from dorsal to ventral depths (dashed trace, R^2^=0.83, p=0.001). Figure 6c reports the session-averaged estimates of relative onset latencies in the visual epoch across depths. For VG trials, visual latencies were detected between –3.6ms (95%CI [-6.2 −1.3]ms) on channel 5 and 9.3ms (95%CI [2 18.8]ms) on channel −8 relative to visual burst onset on the reference channel, and the trend of longer latencies from dorsal to ventral was well captured by the cubic fit (R^2^ =0.66, *P*=0.004). For MG trials, visual latencies were detected between –4.3ms (95%CI [-9.4 0.5]ms) on channel 5 and 7.0ms (95%CI [1.0 12.0]ms) on channel −7 relative to visual burst onset on the reference channel, and the trend of longer latencies from dorsal to ventral was also captured by the cubic fit (R^2^ =0.65, *P*=0.004). Linear regression analysis applied to these data gives similar trends albeit a less good fit for MG trials (VG: R^2^ =0.6, *P*=0.01; MG: R^2^ =0.36, *P*=0.1). As expected, both types of tasks show the same trend in visual burst onset from dorsal to ventral depths within SC, spanning 7.3ms (between channel 5 and channel −8) and 6.8ms (between channel −5 and channel −7) for VG and MG tasks, respectively. This is consistent with single-synaptic transmission from one channel to another in depth, although other mechanisms are also viable (see *Discussion*). A similar pattern in spike timing was recently reported across cortical layers in rodents ^25^.

**Figure 6.**
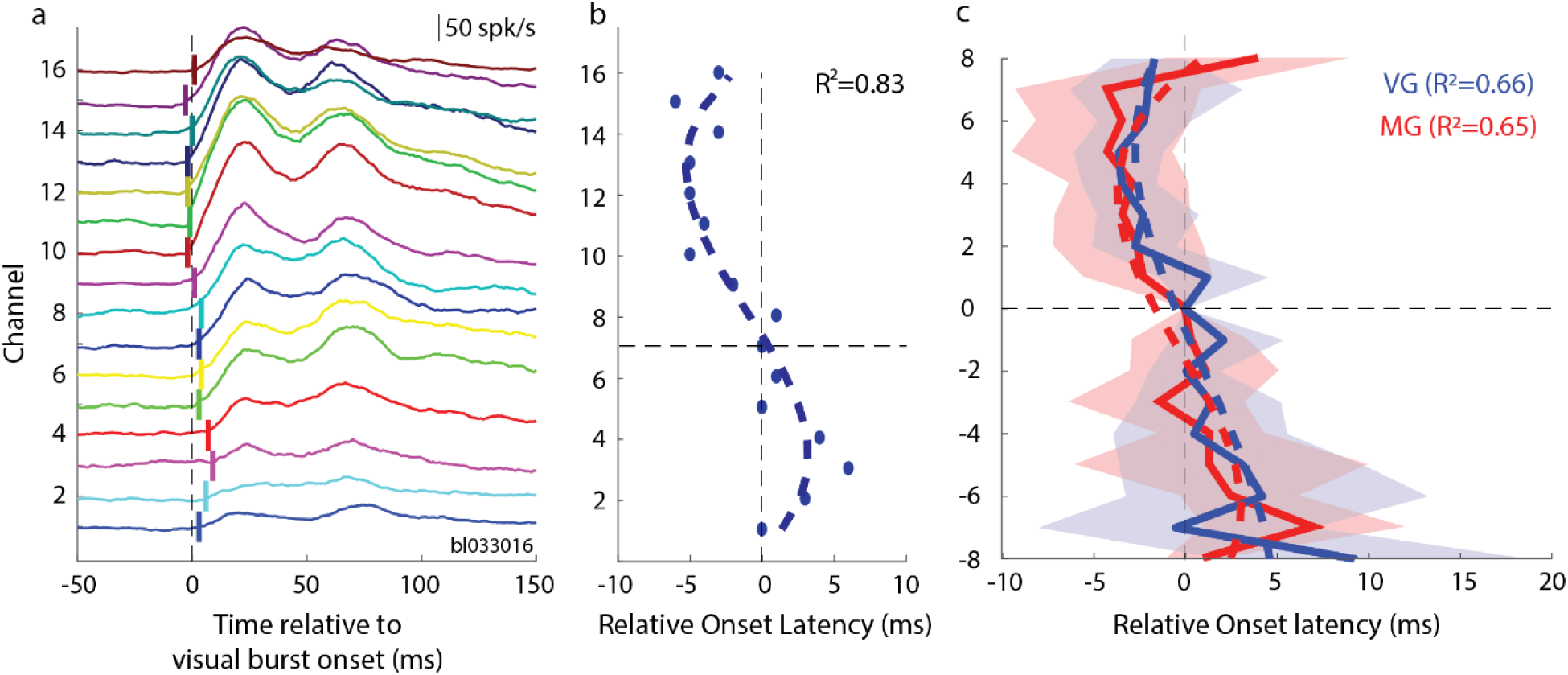
Visual latencies. (a) Data from example VG session showing trial-averaged spike density functions aligned on visual burst onset (see *Materials and Methods*); vertical tick marks indicate the onset of the detected visual activity for each channel; *t* = 0 *ms* corresponds to the onset of the visual activity on the visual alignment channel (here, channel 11). (b) Plot of *relative* visual burst latencies across channels for the same dataset. The data are temporally aligned on the visual onset of the reference channel (here, channel 7), which is required to averaged data across sessions. Dashed trace is a cubic fit (*r*^2^ = 0.84). (c) Session-averaged relative visual onset latencies across depths for VG (thick blue line) and MG (thick red line) trials, respectively. The dashed traces are cubic fits applied separately to the VG (blue dashed trace; *R*^2^ = 0.66) and MG distributions (red dashed trace; *R*^2^ = 0.65). The blue and red translucid regions surrounding the average latency estimations represent the 95% confidence interval computed across sessions; note that the CI of the reference channel is 0 because the activity of each session is temporally aligned to the latency onset of this channel.

Next, we analyzed the pre-saccadic epoch that leads to movement onset. Previous work has shown that SC neurons display a buildup (or prelude) and/or burst of activity during that epoch ^26,27^. In these previous studies, buildup and burst activity were detected by threshold crossing of the averaged firing rate activity computed over fixed temporal window defined for each event separately. This method was designed with the only objective to detect the presence of these events. Here, we wanted to not only detect them but also reliably estimate their onsets times and associated amplitudes.

We developed an algorithm that, first, estimates the onset latencies of well-defined events in the trial-averaged firing rate during the pre-saccadic epoch and, second, classifies them into interpretable spiking activity such as *buildup* and *burst* (see *Materials and Methods* and Supplementary Figure 3 for a visualization). We first defined three events that can be estimated from the spiking activity: (*E1*) the time of significant increase of the detrended activity compared to baseline; (E2) the hinge point before or at *E1*; (E3) the hinge point between *E2* and the time of peak activity (*P*) (see *Materials and Methods* and Supplementary Figure 3g for the definition of ‘hinge point’). The detection of each event was accompanied by a measure of reliability obtained through a bootstrapping procedure. Events that did not meet a reliability criterion were discarded. As a consequence, anywhere between zero and three events were detected for the average spiking activity of each channel. It is important to note that events *E1, E2* and *E3* capture temporal characteristics of the activity and are interpretation-free in terms of neural circuit mechanism. A subsequent classification procedure enabled the interpretation.

Figure 7a-i shows the detection of events *E1, E2, E3* and *P* for the example dataset. The reliability of the detection of each event was measured by the CIs obtained through bootstrapping (see *Materials and Methods*). One can observe that for many channels CIs were well below the 0.6 threshold, indicating a reliable estimation of all events. Panels f and h also report the slopes and CIs around the hinge points for events *E2* and *E3*. The slopes were not significantly different for most of *E2* events, reflecting the slow accumulation of the activity. In contrast, they were significantly different for *E3*, reflecting a sharp increase of the activity. Figure 7i shows the final estimation of all four events, provided that they were statistically significant (see *Neuronal activity categorization, Materials and Methods*). Even though the search window for *E2* was large (within 100ms before *E1* and a total search window of 300ms), *E2* events were systematically detected within 50ms before E1, reflecting the onset of the accumulation that gives rise to *E1*.

**Figure 7.**
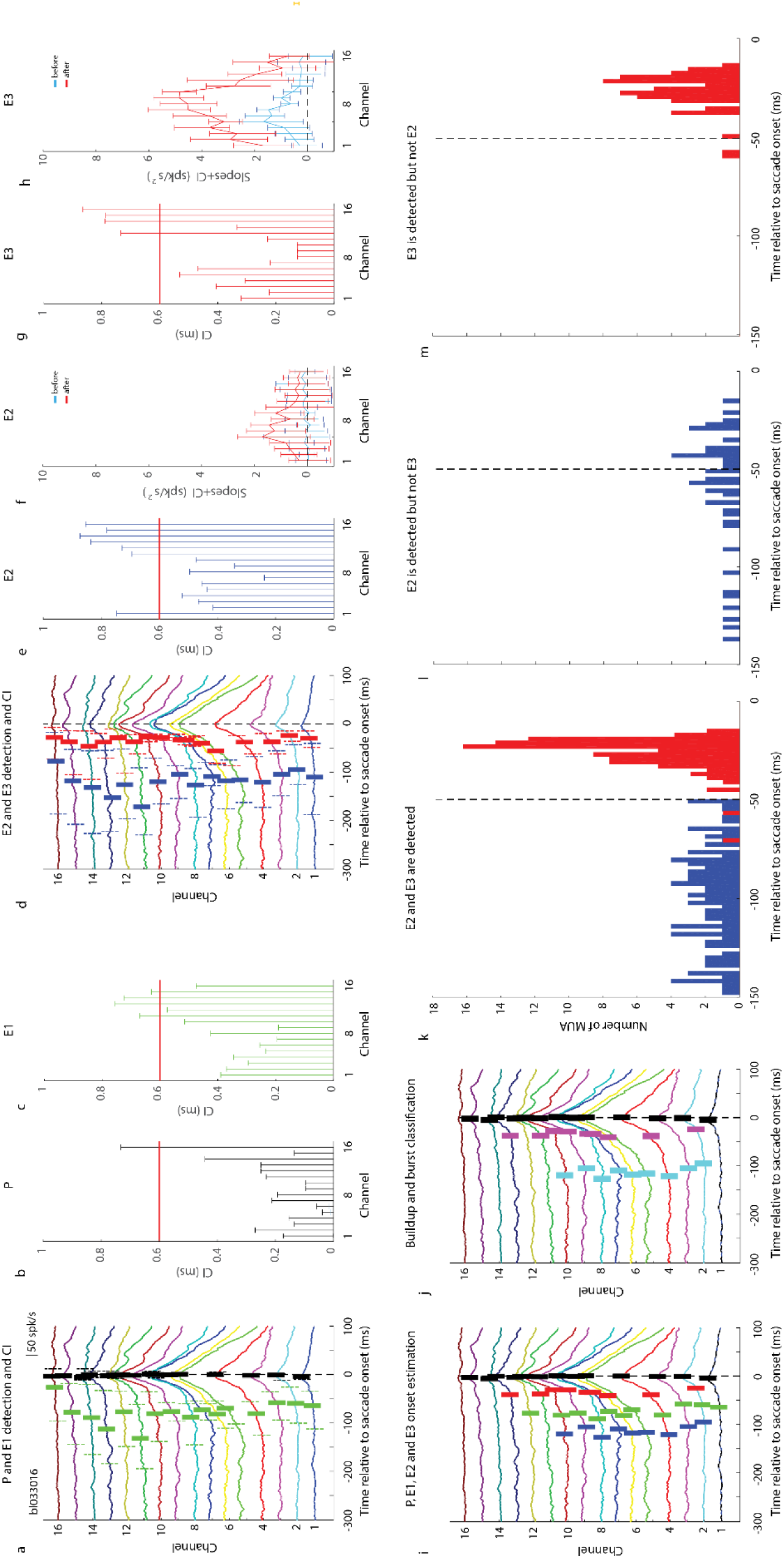
Example of events detection during the pre-saccadic epoch and classification. The detection of all events and their CIs are illustrated for trial-averaged spike density functions on all channels for the VG task of an example session. Data are aligned on saccade onset. Mean event value is shown as a solid, thick vertical tick mark. CIs are denoted as dashed, thin vertical lines of the same color and on each side of the solid tick mark. Events *P, E1, E2* and *E3* are identified respectively in black, green, blue, and red colors. (a-c) Neural activity waveforms are shown with mean and CIs of events *P* and *E1*. (d-h) Neural activity waveforms are shown with mean and CIs of events *E2* and *E3*. (f,h) The mean and CIs of slopes of the linear regressions before (cyan) and after (red) the hinge points are plotted as a function of channel number for events *E2* and *E3*. (i) The spike density waveforms for all channels are now shown with average values of the statistically significant events (see *Materials and Methods* for details). (j) The events are now replaced classification into *buildup* (cyan) or *burst* (purple) events. (k-m) These classifications were determined from the distributions of events *E2* and *E3* and depending on whether only one or both events were detected. We used the −50ms boundary to classify the detected events into *buildup* and *burst* phases of neural activity.

The next step was to classify these events into *buildup* or *burst* activity. When both *E2* and *E3* were detected and were significantly different from each other (i.e., their CIs did not overlap), they were labeled as *buildup* and *burst* events, respectively. If only *E2* was reliably estimated, the activity displayed an early *buildup* (<-50ms) but without a reliable hinge point for *E3* (e.g., channel 3 of Figure 7i). When only *E3* was reliably estimated, the activity only displayed a *burst* just before saccade onset (>-50ms) but without an initial *buildup* (e.g., channels 11 and 13 of Figure 7i). We plotted the distributions of events *E2* and *E3* pooled across all channels and bootstrap iterations, although cases when both or only one of the two events were detected were considered separately (Figure 7k-m). It is important to note that the onset of each event is only constrained temporally by the size of their respective search windows, which means that either onset can occur at any time within 300ms before saccade onset with *buildup* preceding *burst* onset. The distributions of event times are binomial and exhibit visual separation around −50*ms* relative to saccade onset. We therefore used this boundary to distinguish *buildup* (< −50*ms*) from *burst* (> −50*ms*) events. Thus, many events detected as *E2* (*buildup*) were reclassified as *E3* (*burst*). Figure 7j replots the average spike density functions of all channels with identified buildup and burst events superimposed. For this session, peak activity (black tick marks) occurred around saccade onset for nearly all neurons along the dorsoventral dimension, *burst* activity (purple tick marks) was also present in most neurons (but fewer than those with peak activity), while *buildup* activity (cyan tick marks) was present primarily in neurons found along the ventral half of the track.

The analysis shown in Figure 7 was performed for each laminar recording session. We then averaged across the sessions after depth aligning the data relative to the reference channel defined by the CSD method, as discussed earlier (see Supplementary Figure 1). Figure 8 provides a detailed description of the evolution of onset latencies and amplitudes of each event across depths. All results are reported between channels −8 and 8 and 0 was the reference channel. Results are reported separately for VG (Figure 8a-d) and MG (Figure 8e-h) trials.

**Figure 8.**
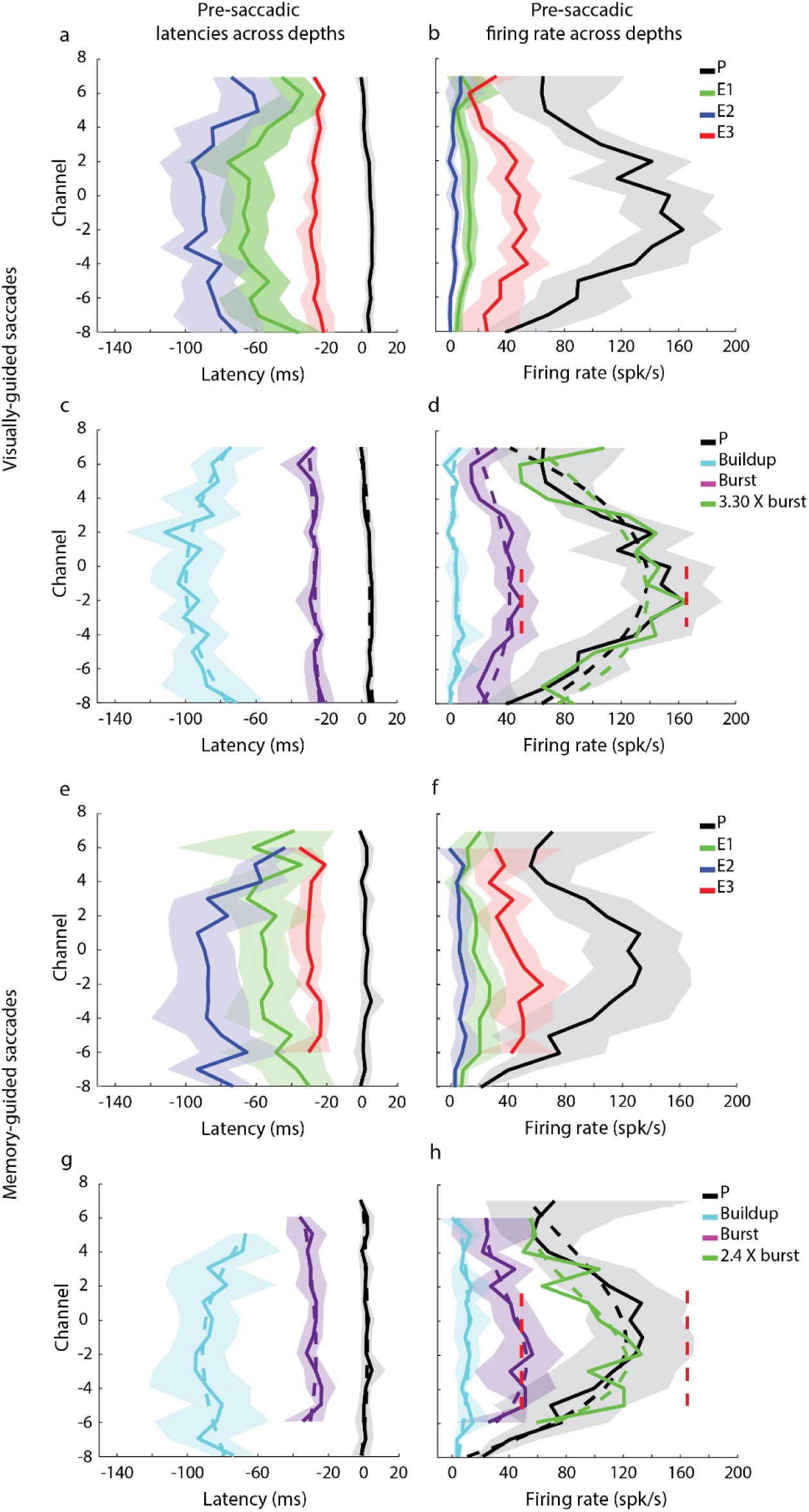
Laminar organization of onset latencies and amplitude of activity during pre-saccadic epoch. (a) Session-averaged onset latencies of events *P* (black trace), *E1* (green trace), *E2* (blue trace) and *E3* (red trace) across depths for VG trials. (b) Session-averaged amplitude of the activity at the time of the events *P, E1, E2* and *E3* across depths for VG trials. (c) Session-averaged onset latencies of *peak* (copy of *P* in panel a, black trace), *buildup* (cyan trace) and *burst* (purple trace) neural activity across depths for VG trials. (d) Session-averaged amplitude of the activity at the time of *peak* (copy of *P* in panel a, black trace), *buildup* (cyan trace) and *burst* (purple trace) onsets across depths for VG trials; a scaled version of the amplitude of *burst* activity relative to *peak* (see *Materials and Methods*) is shown in green; the applied factor is 3.3. (e-h) Data for MG trials are shown following the same format used in panels (a-d) for VG task data. (h) *Burst* activity pattern was multiplied by 2.4 to match the depth-dependent pattern of peak activity. (d,h) Vertical dashed red lines indicate the maximum average activity of *peak* and *burst* for VG trials. (c,d,g,h), dashed lines represent a cubic fit regression. In all panels, the color-matched translucid region surrounding each trace represents the 95% confidence interval computed across sessions.

Figure 8a show the results of the estimation of the events *E1, E2, E3*, and *P*. We found that significant peaks of activity were detected across the whole track (from channel −8 to 7, Figure 8 a,b (black trace)). Event *P* (black trace), which denotes the peak activity, was detected between −0.5ms (95%CI [-4.1 3.5]ms) on channel 7 and 5.9ms (95%CI [3.3 8.4]ms) on channel −2relative to saccade onset. Event *E1* (green trace), which corresponds to a significant change from baseline of the detrended activity during the 200ms preceding saccade onset, was detected between −76.0ms (95%CI [-101.0 −54.2]ms) and −33.8ms (95%CI [-44.6 −24.2]ms) relative to saccade onset. Event *E2* (blue trace), which detects the hinge point associated with the onset of the non-detrended activity accumulation in a 100ms window preceding *E1*, varied between −100.2ms (95%CI [-115.0 −84.6]ms) and −59.0ms (95%CI [-81.7 −35.8]ms) before saccade onset. Figure 8a highlights the tight relationship between *E1* and *E2*, with a maximum average difference of ∼25ms and confidence intervals of the same order for both events, even though *E2* was searched for in a 100ms window preceding *E1*. We interpret this consistency to suggest that the ‘hinge point’ *E2* is a meaningful event. Event *E3* (red trace), identified as the ‘hinge point’ between *E2* and *P*, was detected between −29.2ms (95%CI [-38.0 −23.1]ms) and −21.4ms (95%CI [-22.2 −20.5]ms) before saccade onset.

Figure 8b plots the amplitude of spiking activity at the time of event *P, E1, E2* and *E3* across depths. The activity at the onset of *E1* was limited to low firing rate, between 5.0spk/s(95%CI [2.2 9.4]spk/s) on channel −8 and 16.2spk/s(95%CI [4.0 37.1]spk/s) on channel 6. As expected, the average activity at the onset of *E2* was even closer to baseline than *E1*, and varied between −0.48spk/s (95%CI [-3.4 2.6]spk/s) and 7.8spk/s (95%CI [-3.2 25.5]spk/s). The activity at the onset of *E3* was variable across depths with a maximum amplitude of 54.4spk/s (95%CI [40.3 72.0]spk/s) on channel −4 and the minimum amplitude for the most dorsal and ventral channels. Finally, the activity at event *P* was similar to *E3* but with larger amplitude. Activity at *P* reached a maximum amplitude of 163.3spk/s (95%CI [134.6 192.7]spk/s) on channel −2 and a minimum amplitude for the most dorsal and ventral channels. Overall the results presented in Figure 8a,b indicate that events *E1, E2* and *E3* can be estimated reliably in the average waveform on each channel and that they correspond to systematic events of the pre-saccadic activity.

Once the events (*E2, E3*) were detected, we classified them into *buildup* and *burst* categories. Figure 8c (cyan traces) shows the organization of the onset of *buildup* events across depths. Onset of *buildup* was the earliest on channel 2 with an average latency of −102.7ms (95%CI [-120.2 −83.1]ms) relative to saccade onset. It occurred gradually later for increasingly more distant channels in both dorsal and ventral directions, reaching a minimum of −71.1ms (95%CI [-97.2 −55.2]ms) relative to saccade onset. These onsets estimates are similar albeit slightly later than estimations obtained by the previous study which analyzed the onset of *buildup* 100ms before saccade onset ^26^. The difference in technique to extract onset of *buildup* may explain this discrepancy. Also, our capacity to record across depths with a laminar probe allowed a finer sampling of the neural activity than this previous study that focused mainly on large neural activity. Interestingly, the onsets of *buildup* activities displayed a systematic shift across depths and was well captured by a cubic fit (dashed cyan line, R^2^=0.67, *P*=0.003) reaching a minimum on channel −1. While this analysis cannot reveal a causal relationship between the activity of neurons on different channels, the gradual change of the onset latency of *buildup* may be the result of an activity that initiated in neurons situated around channel −1 and progressively later in dorsal and ventral neurons. This spatiotemporal pattern across depths may reflect a network process that leads to the generation of the movement-related burst (see *Discussion*). Figure 8d (cyan trace) shows that the firing rate at the onset of the *buildup* activity was on average 3.4spk/s (95%CI [-0.3 8.3]spk/s) without any significant trend across channels (cubic fit, R^2^=0.31, *P*=0.21) indicating that the detection of the onset of *buildup* corresponded to a true onset of activity relative to baseline.

The purple trace in Figure 8c shows the organization of *burst* onset across depths. *Bursts* were found at all recorded depths, from channel −8 to channel 7. All *burst*s appeared on average −26.9ms (95%CI [-32.0 −22.3]ms) relative to saccade onset which is similar to a previous result (Figure 18 in ^26^). All *bursts* were temporally tightly aligned (i.e. synchronous, without any significant trend across depths) (cubic fit, R^2^=0.4, *P*=0.11), regardless of whether the *burst* was preceded by a *buildup*. This result is indicative of a general recruitment into a ‘burst’ mode of all neurons along the dorsoventral axis. Figure 8d (purple traces) shows that the firing rate at the onset of the *burst* activity reached a maximum of 50.2spk/s (95%CI [39.9 61.4]spk/s) on channel −2 and decreased for dorsal and ventral channels. This shift was well captured by a cubic fit (dashed purple line, R^2^=0.60, *P*=0.01). Hence, *burst* amplitude displayed a systematic shift across depths: neurons situated on channel just dorsal to the reference channel displayed the maximum firing rate, while neurons at the most dorsal and ventral positions displayed a reduced firing rate. This result implies that the *burst* activity, which is related to the signal that is sent to downstream structures to control the eye movement generation, is not simply duplicated across depths but, on the contrary, its amplitude is a function of the laminar position from where it originates (see *Discussion*).

We repeated the same analyses for MG trials (Figure 8e-h). Figure 8e show the results of the estimation of the events *E1, E2, E3*, and *P*. We found that significant peaks of activity were detected across the whole track (from channel −8 to 7, Figure 8e,f (black trace)). Event *P* (black trace) was detected on average 1.3ms (95%CI [-2.0 4.8]ms) after saccade onset without any significant trend across channels (cubic fit, R^2^ =0.33, *P*=0.2 >0.05). Event *E1* (green trace) was detected between −65.1ms (95%CI [-88.1 − 44.5]ms) and −30.2ms (95%CI [-52.4 −15.1]ms) relative to saccade onset. Event *E2* (blue trace) varied between −93.6ms (95%CI [-110.9 −76.3]ms) and −43.9ms (95%CI [-51.4 −36.5]ms) relative to saccade onset. Once again, Figure 8e highlights the tight relationship between *E1* and *E2*, with a maximum difference of ∼30ms and confidence intervals of the same order for both events. Event *E3* (red trace) was detected on average −28.4ms (95%CI [-36.4 −22.1] ms) relative to saccade onset between channel −6 and 6, without any significant trend across depths (cubic fit, R^2^=0.10, *P*=0.80).

Figure 8f plots the amplitude of spiking activity at the time of event *P, E1, E2* and *E3* across depths. The activity at the onset of *E1* was limited to low firing rate between 3.6spk/s(95%CI [-4.7 10.7]spk/s and 27.2spk/s (95%CI [15.6 37.7]spk/s. As expected, the average activity at the onset of *E2* was very close to baseline, between −1.1spk/s (95%CI [-10.6 9.2]spk/s) and 11.3spk/s (95%CI [4.4 18.0]spk/s). The activity at the onset of *E3* was variable across depths with a maximum amplitude of 63.8spk/s (95%CI [43.7 82.8]spk/s) on channel −2. The minimum amplitude for the most dorsal channel 26.9spk/s (95%CI [17.6 37.4]spk/s) on channel 4. Finally, the activity at event *P* reached a maximum amplitude of 132.8spk/s (95%CI [97.1 167.9]spk/s) on channel −1, which is ∼30spk/s lower than the maximum amplitude for VG trials and a minimum activity for the most dorsal and ventral channels. Similar to VG trials, the results presented in Figure 8e,f indicate that events *E1, E2* and *E3* can be estimated reliably in the average waveform.

The classification of events using classification labels (*buildup, burst*) are presented in Figure 8g,h (cyan traces for *buildup* and purple traces for *burst*). Onset of *buildup* was the earliest on channel −3 with an average latency of −95.1ms (95%CI [-119.0 −75.5]ms) relative to saccade onset. It occurred gradually later for increasing more distant channels in both dorsal and ventral directions, reaching a minimum of − 66.7ms (95%CI [-75.0 −59.9]ms) relative to saccade onset. Similar to VG trials, the onsets of *buildup* activities displayed a systematic shift across depths which was well captured by a cubic fit (dashed cyan line, *R*^2^=0.62, *P*= 0.02). Figure 8h (cyan trace) shows that the firing rate at the onset of the *buildup* activity was on average 8.4spk/s (95%CI [1.5 16.0]spk/s) and was not significantly different across channels (cubic fit, R^2^=0.37, *P*=0.14). Similar to VG trials, this indicates that the detection of the onset of *buildup* correspond to a true onset of activity relative to baseline.

The purple trace in Figure 8h shows the organization of *burst* onset across depths. *Bursts* were detected on nearly every neuron encountered in the penetration, from channel −6 to channel 6. Similar to VG trials, all *burst*s appeared on average −29.1ms (95%CI [-36.2 −22.7]ms) relative to saccade onset and all *bursts* along the dorsoventral axis were temporally tightly aligned (i.e. synchronous) (cubic fit, R^2^=0.35, *P*=0.26). Figure 8h (purple traces) shows that the firing rate at the onset of the *burst* activity reached a maximum of 55.7spk/s (95%CI [31.4 78.2]spk/s) on channel −2 and decreased for dorsal and ventral channels. This shift was well captured by a cubic fit (dashed purple line, R^2^=0.70, *P*=0.017). Hence, similar to VG trials, *bursts* amplitude displayed a systematic shift across depths.

Next, we analyzed the correspondence between the activity at *burst* onset and at the peak response. To do so, we used the cubic fit computed on the distribution of the average firing rate at the onset of the *burst* and at the peak *P*. A scaling factor was computed between the maximum firing rates of the two fits. Figure 8d shows the rescaled fit of the *burst*s (green dashed line) for VG trials and shows the close correspondence with the fit of the *peak* across all depths. Hence, the *peak* activity of the movement epoch appears to be a scaled version of the activity at the *burst* onset across depths. Figure 8h shows the same information for MG trials, and shows that the correspondence with the fit of the *peak* is close for ventral channels but noisier for dorsal channels. For VG and MG trials the scaling factor was 3.3 and 2.4, respectively. Note that the amplitude at the onset of *burst* activity for VG and MG trials were not significantly different (Wilcoxon rank sum test, P>0.14). This result reveals that the lower scaling factor between *burst* and the peak activity or MG trials is not the result of higher amplitude of *burst* onset activity. Rather, this indicates a reduced peak activity for MG trials relative to VG trials while burst activity reached a similar amplitude (see *Discussion*).

Finally, we looked at the classification of the spiking activity based on the type of events displayed during the pre-saccadic epoch. Similar to the work of Munoz and Wurtz ^26^, we used three labels based on spiking activity: *buildup*-only, *burst*-only and *buildup*-*burst*. Note that even if their exact definitions are different, buildup and burst activity by the ‘threshold method’ of ^26^ and *buildup* and *burst* events by our technique relate to similar features of the spiking activity and their detection can be used to compare the categorization of SC activity. For VG trials we found that each type of activity represents 17%, 30% and 53% (27%, 29% and 44%, respectively, for MG trials) of the whole population of recorded activity, respectively (Figure 9). Munoz and Wurtz found that *burst-only* neurons was the largest proportion (Table 1, 6%, 68% and 26%, respectively, from ^26^). Our results for both VG and MG trials show that *buildup*-*burst* activity was the largest type. The reason for this discrepancy is two-fold. First, Munoz and Wurtz used a conservative method based on threshold-crossing on the average activity measured in a window starting 100ms before saccade onset. However, the latency distribution of *buildup* onset (Figure 8c) shows that many channels display a *buildup* activity later than 100ms before saccade onset. Hence, their method would not be able to detect these ‘late’ *buildup* onsets. Second, our method is more sensitivity at detecting reliable small *buildup* activity as it is based on the detection of a hinge point of the activity, which is particularly critical for the most dorsal and ventral channels where the average activity is lower. Hence, their method likely yielded a very conservative estimate of the proportion of *buildup* activity. Our laminar data allow us also to plot the distribution of each type of activity across depths (Figure 9). For VG trials, *buildup*-*burst* neurons were found in the more central positions between channels −6 and 4 and less at dorsal positions, *burst*-only neurons are found at all depths but particularly at more dorsal positions above channel 4, and *buildup*-only neurons are found at more ventral positions below channel −7. Similar distribution, albeit noisier, were found for MG trials. These results confirm the subdivision of SC intermediate layers into a dorsal subdivision that mainly contains *burst*-only neurons and a ventral subdivision that mainly contains *buildup*-*burst* neurons ^26^. The boundary of this subdivision would be situated around channel 3.

**Figure 9.**
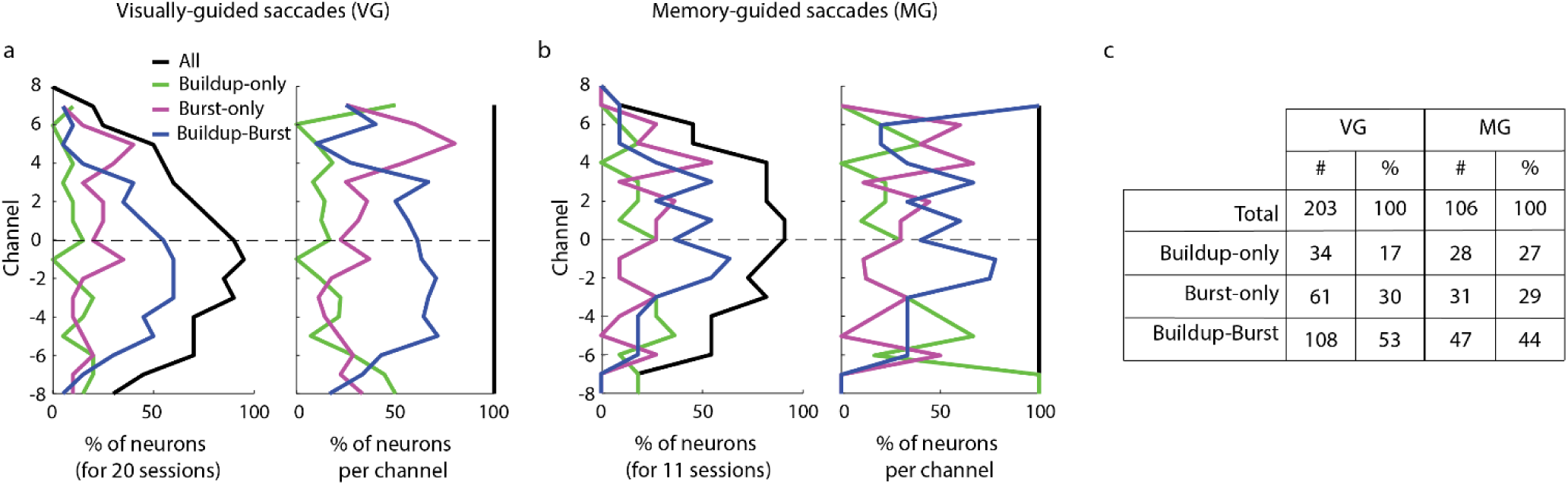
Classification of SC neurons based on pre-saccadic activity. Every panel shows the distributions of buildup-only (green trace), burst-only (purple trace) and buildup-burst (blue trace) neurons as a function of channel number. Data are pooled across 20 VG sessions (a) and 11 MG sessions (b). The figure follows the format of Figure 4. In each subplot, the abscissa denotes the proportion of neurons. Left: For each channel, the neuron count (either for each category or for all neurons) across all sessions is normalized by the number of sessions (20 for VG and 11 for MG trials). Right: The neuron count is normalized individually for each channel to compensate for the non-uniform sampling of neurons across depths (i.e., the neuron count becomes 1 on every channel). (c) A summary of the percentages of each type of neuron for VG and MG trials.

### Stimulation-evoked saccades along the dorsoventral axis of SC

Although the focus of this study was to examine SC activity patterns across depth, we did occasionally deliver electrical stimulation through each contact at the end of experimental sessions. To preserve the integrity of the electrode, however, each contact was stimulated only a few times (typically just twice) and the same stimulation parameters (40µA, 400Hz, 200ms; biphasic, 200µs pulse duration, 17µs inter-pulse duration) were used across contacts and sessions. Sufficient data were available for 12 sessions to permit a preliminary evaluation. Figure 10 plots stimulation-evoked saccade vectors as a function of depth for 5 example sessions. It also provides a histogram of the standard deviations in amplitude and direction across contacts for each session. All but one session had lower than 15° of standard deviation in directions and all sessions had less than 5° of standard deviation in amplitude. Hence, there was a high degree of similarity between saccade vectors across channels. Note that although the electrode was inserted roughly orthogonally to the surface of SC, these measurements are not sufficient to ensure that it traversed in an actual anatomical “column” within the SC.

**Figure 10.**
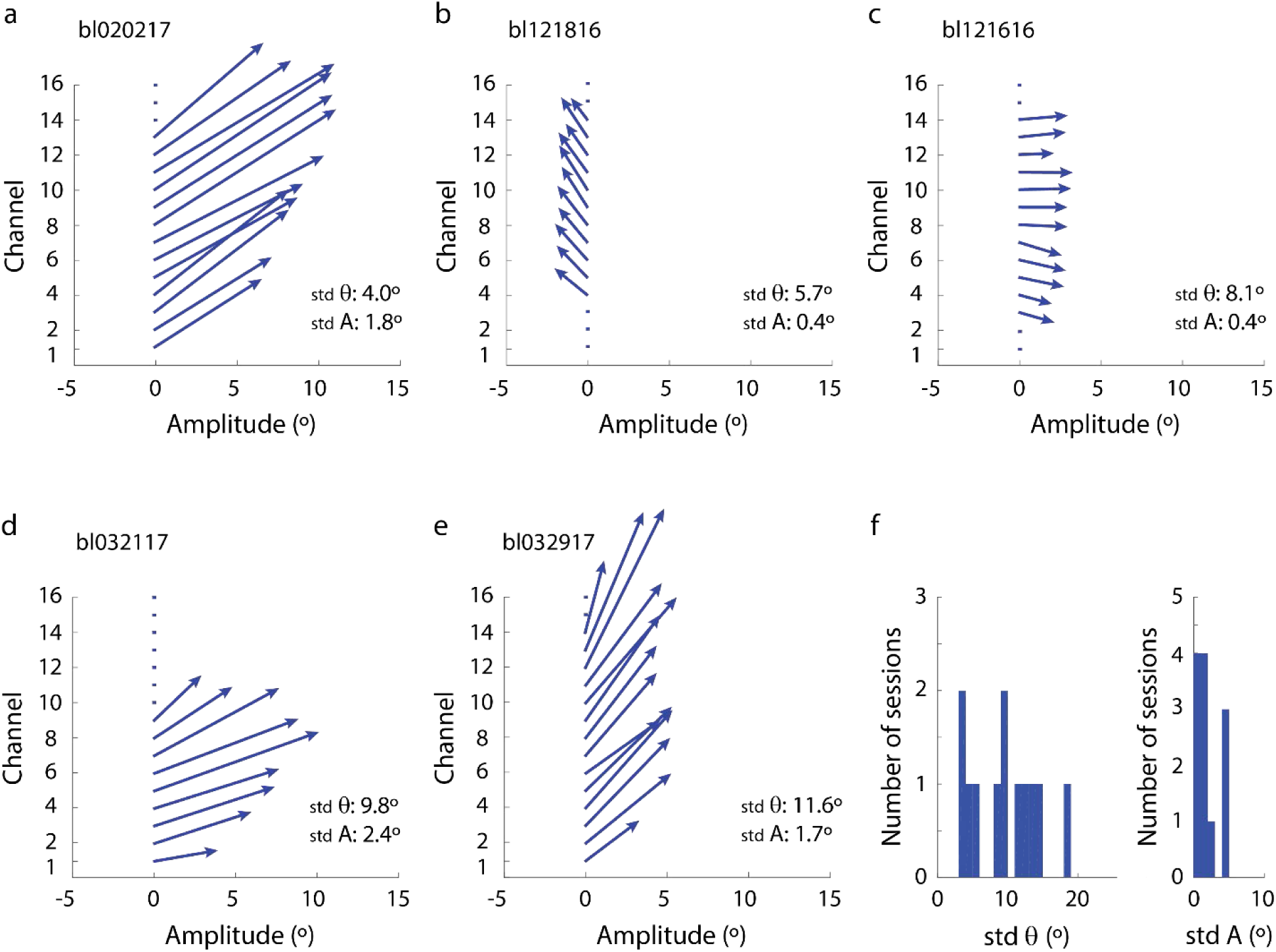
Saccade vector distribution across depths and across sessions. (a-e) Vector of the saccadic eye movement induced by stimulation of each channel during example sessions; blue arrows indicate the direction and amplitude of the induced saccades; blue dots indicate the absence of saccades after the onset of the stimulation; bottom right corner displays the standard deviation of the direction (*θ*) and amplitude (*A*) of the saccade vector for the example session. (f) Distribution of the standard deviation of the direction (left) and amplitude (right) across all sessions when the data from the stimulation paradigm was recorded (12 sessions).

## Discussion

We used a multi-contact laminar probe to record simultaneously the activity of a population of neurons along the dorsoventral axis of the SC of nonhuman primates performing delayed saccade tasks. The categorization of the activity of each channel revealed, as summarized in Figure 11, a visual preference for the most dorsal channels, a movement preference for the most ventral channels, and combined visual and movement responses for intermediate channels. The firing rates associated with the two events were not randomly distributed but rather changed systematically along the dorsoventral dimension (gray shades), each peaking at a certain depth and exhibiting weaker bursts with distance. Maximal activity during the visual epoch was observed ∼1mm more dorsally than during the saccade-related burst, but both were within the intermediate layers. The visuomotor index indicated a clear non-linear relationship along the dorsoventral axis from visual to motor preference. Low-frequency activity observed during the delay period was also not uniform. It followed the same spatial organization as the activity during the visual epoch: the most vigorous visual burst and the strongest delay-period activity were observed at the same depth. The onset latencies of visually-evoked activity revealed a continuous trend from dorsal to ventral channels (purple arrows). The onset latencies of both *buildup* and *burst* activities were detected reliably and revealed systematic spatiotemporal patterns during the pre-saccadic epoch: *buildup* activity was initiated in the central part of the intermediate channels and gradually later in adjacent dorsal and ventral channels (blue arrows), while *burst* activity appeared synchronously across almost all channels (red arrows). These results reveal that SC is functionally organized across depths, and its spatiotemporal patterns reflect network processes properties that were difficult to appreciate in previous studies that relied on single unit recordings. These structural patterns of SC network architecture can inform the design of biologically-inspired models that implement sensorimotor transformation ^6,7^.

**Figure 11.**
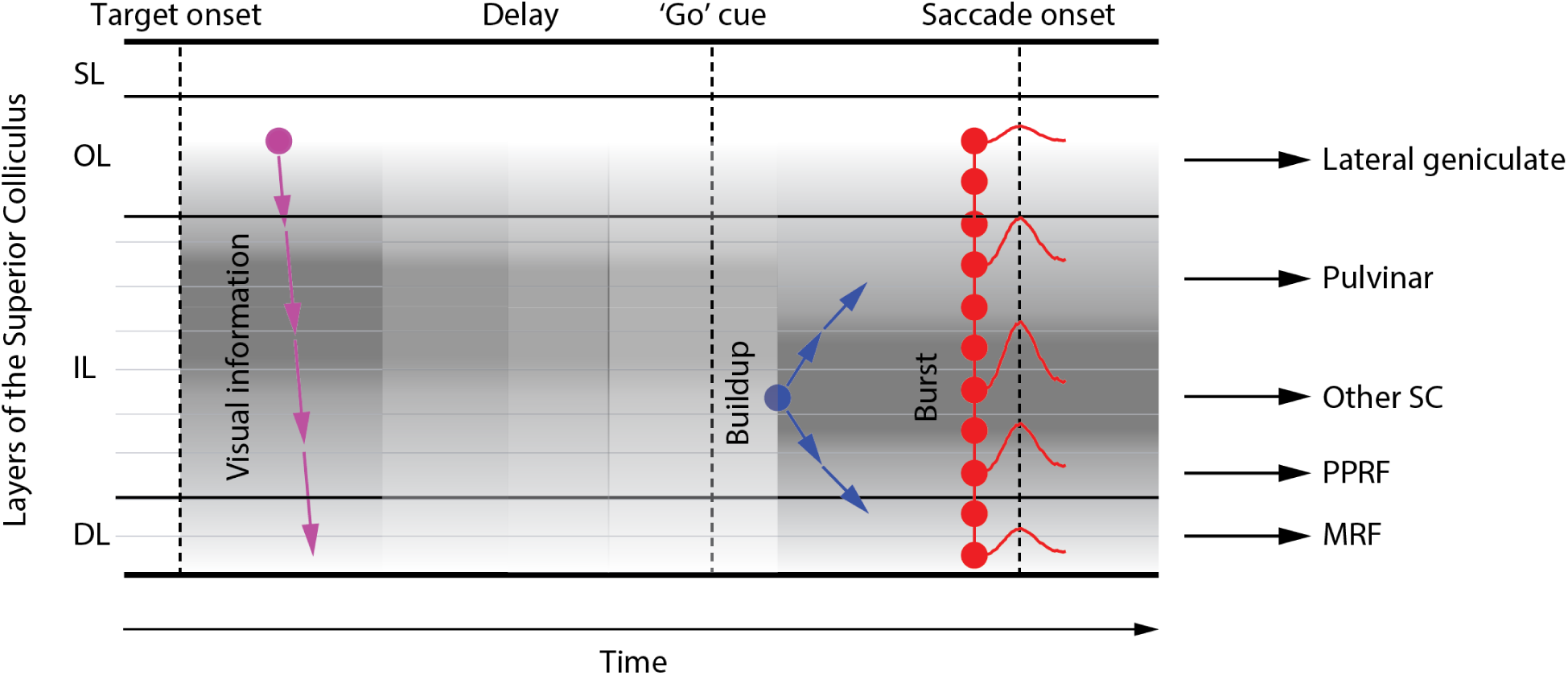
Schematic representation of spatiotemporal dynamics of population activity in SC. Visual information initiates (purple dot) around the superficial and optic layers (SL, OL) and systematically later in sequential order ventrally through the intermediate and deep layers (IL, DL). Neurons in the dorsal intermediate layers produced the most vigorous visual burst, shown as the darkest of the gray shade. The peak firing rate of the visual burst decreased with distance, indicated by lighter shades of gray. In the ensuing delay period, SC neurons exhibit a more sustained low-frequency activity and with a laminar organization that matches that of the visual burst. Approximately 100ms after the ‘go’ cue (blue dot; presumably once fixation offset is processed), neurons around the center of the intermediate layers gradually increase their firing rate. This buildup of activity appears later on adjacent layers both dorsally and ventrally (diverging blue arrows), ultimately leading to a burst synchronously across the entire dorsoventral extent of SC (red dots). Approximately 25-30ms later, a saccade is triggered. The layers with maximal activity during the pre- and peri-saccade periods are shown in gray shades. Note that the SC layer where activity begins to accumulate after the ‘go’ cue is also the layer that is maximally active during the burst, and that neurons maximally active during the movement phase are located more ventrally than neurons maximally active during the visual and delay periods. Rightward horizontal arrows indicate the main outputs form SC and their projection structures (adapted from ^1^).

Past efforts to correlate activity features to neuron location in SC were limited by the uncertainties of estimating the single electrode’s depth. Even the most methodical approach e.g., ^23^ cannot account for settling of neural tissue or of anisotropies in SC geometry (e.g., curvature). The constant intercontact distance of the laminar probe combined with current-source density alignment circumvent many limitations and thereby allow a more rigorous examination of the effects of depth. Consider, for example, the visuomotor index (VMI), a conventional ratio that contrasts the relative contributions of the visually-triggered and movement-related activities of a neuron. The relationship between VMI and depth for single electrode data (see Figure 3B of Ikeda et al. (2015)) shows a linear trend of increasing motor dominance with depth, but it isn’t able to reveal the saturation of VMI at deeper locations that we report here (Figure 3C). This saturation is observed at deeper sites, where the peak activities in the visual and movement epochs begin to decrease in relatively equal amounts. These sites are ventral to the high-frequency, saccade-related burst neurons classically associated with the SC and, we speculate, that sampling bias probably contributed to their omission.

### Insights into sensorimotor transformation

Recognizing that connectivity along the dorsoventral extent of SC is ideally suited to transform sensory signals into movement commands, previous studies have attempted to delineate the functional and anatomical substrates of the transformation. Visual latency, for example, is known to increase only modestly with depth ^28^, on the order of 10-20ms and replicated by our observation (Figure 6C). One logical prediction is that the superficial layers anatomically innervate the intermediate and deep layers of SC and that visual information is relayed through it. Indeed, *in vitro* slice studies in rodents have established this circuit ^4^. However, this pathway is generally considered in the context of converting sensory signal into a movement command and as the primary pathway that governs short-latency express saccades ^29^. While we don’t dispute this hypothesis, we also can’t refute the possibility that this pathway’s central role may be to relay the visual signal to visuomotor neurons. It may even be relayed to putative motor neurons in the SC but other inhibitory inputs may suppress its expression, and the removal of inhibition could unmask the sensory burst ^30^. It is also possible that extracollicular sources may contribute to or augment the visual response relayed from the superficial layers. Indeed, visual responses of visuomotor neurons in intermediate/deep SC are delayed or absent after visual cortex lesion, while the sensory response of visual neurons in superficial layers are not as compromised ^31,32^. Visual information can even be processed through the frontal eye fields, whose projections terminate in the intermediate layers of SC ^33,34^. Future experiments that combine laminar probe recordings with experimental manipulations of extracollicular inputs could provide useful insights into layer-specific functional contributions.

The transient burst of the visual response was followed by low-level activity in a subset of SC neurons (Figure 5). Given the rich balance of excitation and inhibition across all layers of the SC ^35^, the persistent activity can be readily generated through network dynamics e.g., ^36^, although intrinsic, biophysical features likely contribute as well ^37,38^. That this low frequency activity is more prevalent in dorsal layers can be inferred from data compiled with single electrode experiments see Figure 7C,D of ^39^, although its alignment, with rank order preserved, is best appreciated from the laminar probe data (compare Figure 3a,b with Figure 5). Notably, there was no transition from visually-dominant to motor-dominant layers during the delay period, which we believe has major implications on reference-frame transformation research. Previous studies have suggested that the transformation between reference frames occurs during the delay period (under appropriate task design). Correlative evidence exists for craniocentric to oculocentric representation ^40,41^ and for visual, target-centered to motor, gaze-centered coordinates ^42-44^. One way to reconcile these seemingly discrepant results, particularly with respect to the latter set of studies, is that the transformation may occur in dorsal, visually-dominant layers and that the population activity doesn’t transition to the motoric layers until after the animal receives permission to produce a movement. This notion suggests that the gaze-centered signal, although thought to be a movement signal, is in fact not interpreted as a movement command perhaps because the population activity exhibits state space dynamics that are not optimal for evoking a movement ^45-47^.

Once the animal receives permission to initiate a saccade, the population activity transitions to more ventral layers (Figures 8 and 11), where neurons begin to accumulate activity ∼100ms before saccade onset. Slice experiments suggest that *buildup* activity is mediated by both a reduction of GABAergic inhibition ^29^ and amplification by NMDA-mediated synaptic transmission in local excitatory circuitry within the intermediate layer neurons ^48-51^. This local excitatory circuit, perhaps along with the excitatory ascending pathway ^52,53^, induces *buildup* activity in neighboring neurons in adjacent layers at gradually longer latencies (blue arrows, Figure 11). The continued amplification of buildup activity culminates in a synchronized burst across nearly all layers of the SC (red arrows, Figure 11), where the peak firing rate of the movement burst appears to be a linear amplification of the cell’s activity at *burst* onset ^49^. Accordingly, across the active population, the neurons with the earliest *buildup* onset accumulate activity the longest and therefore have the highest firing rate at both *burst* onset and at peak. Given their saccade related discharge profiles, these are the putative neurons that project to the brainstem burst generator ^54,55^ and likely mediate instantaneous control of saccade velocity ^56,57^. Further, the constant scaling factor between activity at burst onset to peak burst across channels (Figure 8) provides functional evidence of linear amplification in the motor burst ^49^. The amplification factor was different for VG and MG trials (∼3 vs. ∼2) while the activity at *burst* onset was similar. Whether SC can realize this computation intrinsically remains an open question and will be the object of future research.

Finally, as suggested in Figure 11, each laminar position within SC contains projection neurons to different structures. The laminar organization of the movement burst amplitude implies that these projection structures may decode the output of SC information in a specific way, maybe reflecting different constraints related to their role in the control of the eye movement generation. For example, the signal sent to PPRF may need to be temporally precise while a corollary discharge of the saccade command to pulvinar or relayed to FEF, may not require such precision. Further computational modelling and multi-area recording are required to evaluate the relation between the laminar organization of the SC activity and the input signals to its projection structures.

### Nomenclature of SC neurons

The long history of SC studies of sensorimotor transformation, specifically visual input leading to saccadic eye movement, have yielded a variety of names to describe the types of neurons involved in the process. The most generic, hypothesis-free nomenclature is to classify them as visual, motor and visuomotor neurons based on activity modulation during the visual and/or movement intervals. Visual neurons are found in the superficial layers, and visuomotor and motor neurons reside in the deeper layers 58,59, but it has been debated whether the latter two are segregated along the dorsoventral axis. The ability of a laminar probe to record simultaneously the activities of neurons along this axis revealed that there is a gradual transition from visual to visuomotor to motor neurons with depth but the vast majority are visuomotor (Figure 4). We also demonstrated previously ^30^ and discussed above that putative motor neurons can exhibit a visual response under certain conditions. Thus, we prefer to avoid making a distinction between visuomotor and motor neurons.

Hypothesis-guided names have also been assigned to SC neurons, with different nomenclatures being introduced over time. Saccade-related burst neurons discharge a high-frequency burst for optimal vector saccades, and burst onset is tightly coupled to saccade onset, leading the movement by ∼20ms ^58^. Neurons with low-frequency activity several hundred milliseconds before the burst were once called long-lead movement neurons ^59,60^. They may be the same subclass of neurons that were later termed quasi-visual neurons ^20^ and prelude neurons ^27^. Other families of names (clipped, partially clipped, and unclipped neurons or open- and closed-movement field neurons) emerged from experiments that tested whether SC activity controls dynamic motor error ^26,61^. The current nomenclature labels intermediate/deep layer neurons as fixation, *burst*, and *buildup* neurons ^26,62^. Fixation neurons reside in the rostral pole of the SC. They discharge at a tonic rate during fixation, pause during large saccades, and burst during very-small amplitude saccades, including microsaccades ^62,63^. *Buildup* neurons exhibit low-level activity well before saccade onset and therefore are similar to prelude or long-lead movement neurons. *Burst* neurons are closest to the saccade-related burst neurons. Finally, the reader should keep in mind that despite the use of categories, most neurons exhibit both *burst* and *buildup* features (Figure 9).

More recent SC studies have used a stochastic accumulator framework to correlate features of neural activity with saccade reaction time e.g., ^64,65,66^. Fitting the pre-saccadic activity with a two-piecewise linear regression yielded a time of inflection point that is correlated with reaction time ^65^. Moreover, their data suggest that the accumulation occurs ∼65ms before saccade onset (accumulation and saccade onsets were respectively 142±16ms and 207±20ms, relative to fixation offset). We sought to relate this finding to the *buildup* or *burst* features of neural activity, and an initial glance suggests that accumulation onset is a feature of *buildup* neurons. However, our analysis suggests that *buildup* onset occurs at least 30ms earlier, ∼100ms before saccade onset (Figure 8). We believe this discrepancy may be a result of the differences in detection approaches. We developed a method to reliably detect and classify a *buildup* (accumulation) and/or a *burst* (threshold) process. By contrast, the previous study, by using a two-piece linear regression, limited the detection to only one neural event that spanned from 100 ms before fixation offset to the time of peak in the saccade-related burst ^65^. Their method was applied on individual trials and the amount of stochastic noise in the spiking discharge may have prevented the distinction between two neural events. Here we used trial-averaged waveform combined with bootstrapping of the trial sets, which allowed the detection of at most two distinct neural events within a probabilistic framework. To the best of our knowledge, this is the first time a method is developed to detect precisely and reliably the onset of *buildup* and *burst* activity, beyond the use of predetermined temporal windows of analysis. This method revealed that *buildup* activity was detected gradually on different channels spanning ∼30ms since the initial *buildup* around the center of the intermediate layers.

*Burst* onset occurred synchronously across all layers, ∼28ms before saccade onset. This is comparable with the values used or estimated in previous oculomotor studies ^65,67,68^. It is also in line with the spike modulation times observed in burst generator neurons that participate in saccade generation ^69,70^. It is generally stated the impact of reaching a threshold (equivalently, entering burst mode), either at individual or population activity, is to inhibit the brainstem omnipause neurons ^67,71,72^, although other frameworks that depend on state space dynamical systems may facilitate direct communication between SC and pontine burst neurons ^45-47^.

### Alignment of recorded sessions based on CSD analysis

We presented a method to align SC laminar data across sessions based on the analysis of a sink pattern during the visual epoch. The results show that the alignment is precise enough to unveil a systematic functional organization of the intermediate layers of SC. Here, we interpret the strong sink pattern occurring during the visual epoch as the input current reflecting the visual input from target presentation. Based on known SC neuroanatomy ^1^, this input is most likely occurring in the optic and superficial layers. Previous works in the primary and the frontal cortex ^8,12,13,17,18^ have also used CSD analysis to align data across sessions. Most of these methods use a sink/source pair pattern occurring early on during the trial and interpreted as the sensory input occurring in layer 4. One study, in addition, uses a distinctive pattern of the LFP to measure the depth of the dura while recording in SEF ^18^. In the absence of such anatomical marker, methods based only on a systematic pattern in the CSD, like our SC method or like the cortical methods, can only align data relatively to an arbitrary reference. Histological verification and/or neuroanatomical experiments are required to verify the reference location. We did not pursue this option as the animals are still in use. As next step, it would be valuable to employ CSD profiles to more precisely identify the superficial and deeper boundaries of the SC. To this end, spikes and LFPs would need to be recorded on at least one and ideally more contacts beyond the dorsal and ventral boundaries to evaluate the CSD. Additionally, interpretations will need to account for the different neuronal morphologies observed across layers ^1^ and for the consequences of mechanical damage induced by repeated electrode penetrations across the duration of the experiments.

## Materials & Methods

### Animal preparation

All experimental and surgical procedures were approved by the Institutional Animal Care and Use Committee of the University of Pittsburgh and were in compliance with the U.S. Public Health Service policy on the humane care and use of laboratory animals. Two males, rhesus macaque monkeys (Macaca mulatta, ages 13 and 8 years) served as subjects. Surgical procedure details have been described previously ^73^. Briefly, a recording chamber was placed on each animal to give access to the left and right colliculi. The chamber was tilted 40° posteriorly in the sagittal plane so that probe penetrations were approximately normal to the SC surface. A 3-D printed angular adaptor was also used to adjust the penetration angle to maximize the collinearity of the saccade vectors obtained by stimulation (see below *Neurophysiological recordings*). Head restraint was realized by fitting a thermoplastic mask individually for each animal ^74^.

### Neurophysiological recordings

We used 16-channel laminar probes (Alpha-Omega, Alpharetta, USA; 150µm inter-contact distance; ∼300µm diameter; ∼1M? impedance for each channel) to record neural activity across different layers of the SC (Figure 1a). The probe was advanced with a hydraulic Microdrive (Narishige, Tokyo, Japan). When neural activity was detected on one channel, biphasic electrical stimulation (40µA, 400Hz, 200ms; 200µs pulse duration, 17µs inter-pulse duration) was delivered to the deepest contact to determine its ability to evoke a saccade. Once verified that an intermediate layer of SC was reached, the probe was lowered further until multi-unit activity could be detected on the maximum number of contacts. Electrical stimulation was delivered on different channels to qualitatively gauge that induced saccades had similar characteristics (direction and amplitude) across depths and to estimate the average vector. On a subset of sessions, stimulation was applied systematically on each channel and recorded to provide a quantitative analysis. This vector was used as a measure of the preferred saccade for all units recorded along the different contacts across depths. The saccade endpoint, relative to central fixation, was taken as the location of the center of the visual receptive field. The diametrically opposite location was defined as the anti-receptive field position.

### Data collection

Neurophysiological signals were recorded using the Scout data acquisition system (Ripple, Salt Lake City, USA). Data recording was synchronized with the beginning of the trial, and the timing of all trial events were recorded simultaneously with raw neural activity. For each channel, the raw activity was parsed into spiking activity (high pass filter at 250Hz and threshold at 3.5 times the RMS) and local field potential (low pass filter at 250Hz). As the isolation of single neural activity was not achievable simultaneously on every channel, the recorded spiking activity was always considered as being from multiple units in the vicinity of each contact. A standard threshold crossing was used to determine spike times.

### Behavioral paradigms

After recovery from surgery, each animal was trained to perform standard eye-movements tasks. Eye movements were recorded using an infrared eye-tracker (EyeLink 1000 from SR Research, Ottawa, Canada; RRID:SCR_009602). The camera and the infra-red illuminator were situated vertically above the animal’s head. A hot-mirror was tilted ∼45° between the animal’s head and the display monitor. EyeLink software sampled the pupil at 1KHz in the reflected infra-red image, using its center as gaze position. Calibration of gaze position was performed for each session by having the animal fixate targets displayed at known locations on the monitor. A real-time system was used for the control of the behavioral tasks and data acquisition ^75^. Animals were trained to perform two standard eye movement tasks (Figure 1b): a visually-guided delayed saccade task (VG) and a memory-guided delayed saccade task (MG). At the beginning of a VG trial, a fixation dot appeared at the center of the screen. The animal had 3000ms to bring its line of sight towards it and maintain its fixation for 200 to 350ms. Then a second dot (target) appeared at a specific location on the screen. The animal had to keep its line of sight on the fixation dot for an additional 600 to 900ms. At the end of this delay period, the fixation dot disappeared (“go” cue) and the animal had to make a saccade toward the target within 500ms. The animal had to fixate the target for at least 250ms in order to get a liquid reward and for the trial to be successful. If one of these conditions was not respected, the trial was aborted. The timeline for an MG trial was the same as for a VG trial except that the saccade target remained illuminated only for a fixed period of time (300ms). The animal was then forced to maintain the target position in memory for the remaining of the trial. The two types of trial were typically randomly interleaved. Target locations were randomly interleaved between two locations: the center of the receptive field of the neural activity and the diametrically opposite location (see above *Neurophysiological recordings*). We only present data from trials with the target in the response field.

### Neural activity analysis

During the delayed saccade tasks, neural activity typically increases or bursts shortly after stimulus presentation in the receptive filed and/or during a saccade directed to that target. In between the two bursts, many SC neurons exhibit a low-frequency discharge see Figure 1 of ref. ^21^. We therefore analyzed activity separately for the visual epoch that follows target onset, the delay period that follows the visual burst and continues until fixation point is extinguished, and the movement-related epoch that ensues the ‘go’ cue. Neural activity analyses were performed either on discrete spike trains or continuous spike density waveforms obtained by convolving the spikes with a kernel simulating an excitatory post-synaptic potential (growth and decay times constants of 1ms and 20ms, respectively)^76^. All data analyses were performed in MATLAB (Mathworks, Natick, USA; RRID: SCR_001622).

#### Visual activity

The study of neural response to a stimulus is traditionally performed by aligning the data on target presentation. This approach effectively ignores the trial-to-trial variability in both the actual time of target presentation (due to frame rate limit) and neuronal stochasticity. We were concerned that this variance may be greater than the time spanned for the visual response to emerge across layers. Therefore, we opted to analyze activity during the visual epoch by aligning the data on visual burst onset. We applied on the spike train for each trial and channel the ‘Poisson surprise’ method ^24^ to detect burst onset in the epoch [30 150]ms following target onset. Burst detection criteria were set to a minimum of 3 spikes and a surprise index of –*log* (0.025). For each session, the channel with the maximum number of trials with detected bursts was selected as the *alignment channel*. Trials for which no burst was detected on the alignment channel were discarded. For every remaining trial, the visual burst on the alignment channel was used to align the activity of all the other channels. Note that the activity on all the other channels was used regardless of whether a burst was detected on them. The realigned spike train for each channel was converted to a spike density function and then averaged across trials.

We next determined the relative burst onset times across the channels (Supplementary Figure 2). First, each trial-averaged spike density waveform was baseline-corrected by the mean activity computed on the same channel in the [-150 −50]ms epoch relative to visual burst onset. Next, the time of peak visual activity (*Pv*) was detected in the epoch [-50 150]ms around the visual burst onset. We then performed a procedure to statistically compare the distribution of activity in two sliding 20ms-windows *W1* and *W2* (Supplementary Figure 2a). The number of points in each distribution was equal to the number of time points in the window, and *W2* was always shifted 10ms earlier in time relative to *W1*. Initially, *W1* corresponded to the window [*Pv*-20 *Pv*]ms, and *W2* to the window [*Pv*-30 *Pv-10*]ms. Both windows slid to earlier times in 1ms steps until *W2* reached the window [-50 −30]ms (i.e. the beginning of the visual epoch). For each instantiation, the statistical difference between the distributions of activity in *W1* and *W2* was measured using a *t*-test (p<0.01). Because both *W1* and *W2* started around the peak activity, their distributions were significantly different initially. Sliding to earlier times, the first instance (i.e. the beginning of *W1*) when their distributions became not significantly different and stayed not significant for the next 10 iterations defined *Bv*, the first time point when the activity was not significantly different from baseline (Supplementary Figure 2b). This method was designed to account for the presence of a secondary peak in the visual epoch. Finally, a two-piecewise linear regression was computed to estimate the time point (*Lv*) when the spike density waveform started increasing towards *Pv* (Supplementary Figure 2c,d). The regression analysis identified the intervals [*Bv Lv*] and [*Lv Pv*], respectively, where *Lv* minimized the sum of the residuals of the two linear fits and represented our estimate of the visual latency.

#### Delay-period activity

To examine the distribution of delay period activity across the dorsoventral axis, we plotted average, baseline-corrected activity in 50ms nonoverlapping bins on each channel. The bins spanned from the time of the last peak of the visual burst (there were often two) to the end of the shortest delay period. The method typically yielded 5 to 6 bins for analysis. Baseline activity was the same used for visual epoch analysis.

#### Movement-related activity

Trial-averaged spike density waveforms were aligned on saccade onset, which was detected using a velocity criterion (30deg/s) applied after ‘go’ cue. Each signal was also corrected for baseline activity, defined as the average activity in the epoch [-100 0]ms relative to ‘go’ cue. The movement-related activity was separated into two periods. The peri-saccadic or movement-related burst was quantified as the baseline-corrected average activity in the epoch [-25 25]ms centered on saccade onset. This parameter was used for analyses that computed activity levels during the burst and the visuo-motor index (see below). We also defined a pre-saccadic epoch that started at the go cue and continued until saccade onset. Neural activity in this window was analyzed to detect the presence and the onset of *buildup* and *burst* in activity (Supplementary Figure 3). In terms of stochastic accumulator framework e.g., ^65^, this is equivalent to detecting when the activity begins to accumulate, while accounting for trends induced by baseline activity, and when it transitions into a burst. The objective was to detect up to three events in this period: *E1*, corresponding to the time of significant change of activity compared to baseline (which may include a linear trend, see below); *E2*, denoting when activity begins to accumulate and corresponding to a ‘hinge’ point prior to *E1*; and *E3*, marking when the activity starts to burst and corresponding to a ‘hinge’ point occurring between *E2* and *P*, the time of peak activity around saccade onset. To be able to flexibly detect one and/or two separate events (and subsequently onsets of *buildup* and/or *burst* activity), the Poisson surprise method used before for the estimation of the visual onset latency was discarded in favor of a 2-piecewise linear regression-based approach. We sought to limit the number of ad-hoc parameters while relying on statistical measures of significance through data bootstrapping. Also, this analysis was limited to saccades produced in the standard latency range of 200 to 400ms.

As an initial step in our analysis, we applied a detrending procedure to remove any potential bias contributed by the low-frequency discharge from the delay period, well before the buildup and/or burst processes are engaged. A linear trend was estimated between in the epoch [300 200]ms before saccade onset. The obtained linear trend was extrapolated to the remaining time points and subtracted from the trial-averaged activity. Note that this step was only temporary and that all event detections after event *E1b* (see below) were performed on the raw, non-detrended data.

Event *E1* was defined as the first time-point starting 200ms before saccade onset for which the activity became and remained significantly different from baseline for at least 100ms. Baseline was taken as the distribution of activity during the 100ms period preceding the ‘go’ cue on each trial. The number of points in the baseline distribution was equal to the number of time points in the window multiplied by the number of trials. Statistically significant difference was measured using a *t*-test between the distribution of baseline activity and the distribution of activity across trials at each 1ms time bin (p<0.01). To obtain a robust estimate of *E1* and to measure confidence intervals, subsets of trials were created through bootstrapping. An estimate of *E1* (denoted *E1b*) was obtained for each bootstrap iteration. Supplementary Figure 3a provides a visualization of the method for one bootstrap. We used 100 bootstrapped estimates (Supplementary Figure 3c) and defined *E1* as the average of all 100 *E1b* (Supplementary Figure 3e, left). Confidence intervals (CIs) were used as a measure of the reliability of the estimation of *E1* and computed as the 95% quantile of the *E1b* distribution (Supplementary Figure 3e, right). For all events, CIs were normalized with the average size of the search window. For *E1b*, the size of the search window was constant at 200ms. To exclude unreliable estimation of *E1*, a 0.6 threshold was applied on the total range of CIs. This threshold was the only ad-hoc parameter in the algorithm and the same value was used for all events (which was allowed by the normalization of the CIs). The threshold value was chosen in order to remove very unreliable estimations (for example for event *E1*, the excluded estimations had CIs superior to 120ms). Changing the threshold value (e.g. to 0.5 or 0.4) did not alter the general trend of the results (data not shown). With this method, *E1* is the latest time point of statistically significant change from baseline activity. The actual change, indicating the onset of accumulation, most likely occurs prior to it. Thus, we next operated on the non-detrended averaged spike density waveform to obtain a better estimate of the actual time of change from baseline activity, while imposing the constraint of *E1*.

Event *E2* denotes the time point before *E1* when the activity starts deviating from the ongoing activity and displays what we refer to a ‘hinge point’, which we define as the time point at which the rate of change of the spiking activity deviates from its current trend (see Supplementary Figure 3g for a general visualization). The hinge point was detected by finding the best piecewise linear regression for the relevant data points. For each bootstrapped estimation *E1b*, a piecewise linear regression was performed on the intervals [*E1b*-100 *Hb*] and [*Hb E1b*], respectively, and the value of *Hb* that minimized the sum of the residuals of the total fit was the estimate of *E2b*. The combination of the search windows of *E1b* and of *E2b* relative to *E1b*, implies that *E2b* was searched in a potentially very large window starting 300ms before saccade onset and ending as late as *P*. By definition, *E2b* always preceded *E1b* or was equal to it. *E2* was taken as the average of all the bootstrapped estimates of *E2b*. As before, we performed 100 bootstrapped estimations and computed the CIs, which were normalized by 100ms. To exclude an unreliable estimate of *E2*, a 0.6 threshold was applied on the total range of the normalized CIs (Supplementary Figure 3e,f).

Event *E3* marks when the activity starts to burst and corresponds to the hinge point occurring between *E2* and *P*, the time of peak activity around saccade onset. For each round of bootstrapping, we obtain E2b as stated above and an estimate of the time of peak activity (*Pb*). Then we estimated the hinge point *E3b* by fitting a two-piecewise linear regression between *E2b* and *Pb*. The detection of the slopes of the two linear regressions around *E3b* are stored for statistical analysis. By definition, *E3b* is always between *E2b* and *Pb*. *E3* was taken of the average of all the bootstrapped estimates *E3b*. To exclude an unreliable estimate of *E3*, a 0.6 threshold was applied on the total range of the CIs, which normalized by the average interval between *E2b* and *Pb*. *E3* was considered a hinge point only if the CIs of the distribution of the slopes of the two linear regressions before and after all *E3b* were not overlapping (this is a conservative measure of significance). Hence, all estimated *E3* values correspond to a hinge point with a significant change of rate of activity around this time (Supplementary Figure 3d,f,g). Also, *P* was taken as the average of all *Pb* and confidence intervals were computed (Supplementary Figure 3c,e).

#### Classification of events into *buildup* and *burst* activity

Once estimated, these events were used to categorize the pre-saccadic discharge pattern into *buildup* or *burst* activity (Supplementary Figure 3h). To avoid any confusion with previous literature ^26,27^, we used the terms *buildup* and *burst* with an italic typography to refer to these events with a definition specific to this study. We started with the subset of channels across all sessions for which both *E2* and *E3* were both detected and were significantly different from each other (i.e., their confidence intervals did not overlap). A distribution of the event times exhibited visual separation around −50ms relative to saccade onset (data shown in Results in Figure7k,l,m). We therefore used this boundary to distinguish *buildup* (<-50ms) from *burst* (>-50ms) events. We similarly examined the distribution of times when only one of the two events was detected, and we once again used the −50ms boundary criterion to classify the event. Thus, it was possible that activity associated with event *E2* (*E3*) in the absence of *E3* (*E2*) to be classified as a burst (buildup) activity.

### Neuronal activity categorization

Existing literature uses nomenclature for categorizing the cell types in the SC (and other structures, such as FEF) depending on their significant activity during the visual or the peri-saccadic movement epoch. We also performed analyses to determine how neurons based on this classification vary with depth. Only channels with statistically significant neural activity in at least one of the two intervals were included in the subsequent analyses. To measure the significance of the activity during the visual epoch, we used spike density data aligned on visual burst onset but not baseline-corrected. The significance was measured using a statistical test carried out between the distribution across trials of the activity during the visual epoch and the activity during baseline (Wilcoxon rank sum test, P<0.001). To discard very low activity after baseline correction, an additional low threshold (10spk/s) was used on the trial-averaged baseline-corrected activity and averaged across the visual epoch. We used the same procedure to measure the statistical significance of the activity during the movement epoch with data aligned on saccade onset and baseline activity measured on data aligned on ‘go’ cue (see above). Based on this measure of significance of activity in each epoch, the MUA on each channel was categorized as follows: visual-only activity (significant visual activity and not significant movement activity); visuo-movement activity (significant visual and movement activity); movement-only activity (not significant visual activity and significant movement activity).

The visuo-movement index (VMI) contrasts the visual and the movement activity of multi-unit activity (MUA) during the delayed saccade tasks. The visual activity (*V*) is the baseline-corrected average activity in the epoch [0 100]ms following visual burst onset (see above *Neural data analyses*). The movement activity (*M*) is the baseline-corrected average activity in the peri-saccadic epoch [-25 25]ms centered on saccade onset. We defined the index as *VMI* = (M - *V*)/(M + *V*). With this formulation, *VMI* =−1 corresponds to a visual neuron with no saccade related activity while *VMI*= +1 corresponds to a movement neuron with no visual response. We also computed VMI trends when average activity in each epoch is replaced with peak activity and when baseline activity for the movement epoch is measured before target onset. VMI results were qualitatively similar for these variations (data not shown).

### Depth alignment of multiple sessions

The above analyses focus on population neural activity across SC layers within a session. To assess reliability, data must be averaged across sessions, which required appropriate alignment of data collected in each penetration. To address this issue, we designed a method based on current-source density analysis (CSD) ^77^, which computes the second spatial derivative of LFPs and provides an estimate of the distribution of the current sinks and sources as a function of space and time in a volume of tissue. We estimated the CSD using the *csdplotter* toolbox that contains the implementation of the iCSD method (https://github.com/espenhgn/CSDplotter) ^78^ Supplementary Figure 1 demonstrates the utility of the CSD method for the depth alignment of two datasets recorded at two different locations 1mm apart along the same penetration. Panels a and d display average LFP signals recorded at each contact at two locations 1mm apart along the same penetration. The LFPs showed a large decrease reflecting the input current following the display of the target. The CSD plots of the two datasets (panels b and e) revealed a strong current sink (orange bands) occurring after target onset and which encompassed almost 1mm of SC tissue. This feature was present in the recordings at both locations but translated in depth. The lower bound of the sink pattern was at 1.28mm and 1.87mm in panels b and e, respectively. Such a strong sink appeared systematically after target onset across all recording sites (data not shown here, but subject to a future manuscript). We exploited this feature to align the data across sessions. The lower bound limit of the sink pattern was used as a reference for estimating the relative depth of the probe. Supplementary Figure 1c,f shows the average profile of the CSD in a 150ms window starting before burst onset. The transitions from negative to positive CSD was detected automatically and visually inspected to account for the rare cases when the sink pattern was not continuous due to decreased SNR of the LFP. To assess the utility of the CSD alignment method, we compared the relationship between depth and visuo-motor index (VMI, see below) at the two depths (panel g). The two VMI plots appear very similar but shifted in depth. After alignment using the CSD method, the two VMI graphs overlap very well (panel h).

For aligning data by depth across recording sessions, the channel closest to the transition from negative to positive CSD was identified. We termed it the reference channel and assigned it index 0 in plots presenting data after alignment. The indices of the remaining channels were shifted accordingly. Note that the alignment was done in terms of channel index and not in actual mm, which would require an interpolation of the signals between the channels. Given that the inter-contact distance of the probe is 150µm, the maximum error of alignment based on channel index is 75µm. Supplementary Figure 1i shows the average CSD profile of all sessions and for both VG and MG trials. It highlights the robustness of the detection of the CSD reference channel and the systematic presence of a sink pattern above it (i.e., negative values of the CSD profile). Note that if depth alignment between datasets is necessary (i.e. the probe’s depth position is different across datasets), then there exists at least one non-overlapping channel between them. Thus, if any of these non-overlapping channels contains significant activity, then the total number of channels across which data can be analyzed may be larger than the total number of channels of the probe. In practice, only few channels were added and all data analysis that required depth alignment will be presented between channels −8 and 8 (17 channels).

### Statistical analysis

Trial-averaged activity was computed using large numbers of trials. Confidence intervals were measured using bootstrapping (1000 bootstraps, *bootci* in MATLAB). Normality assumption was systematically assessed using a Kolomogorov-Smirnov test (*kstest* in MATLAB). When the hypothesis of normality was not rejected, a parametric test was applied (t-test, *ttest* in MATLAB) otherwise a non-parametric test was used (Wilcoxon rank sum test, *ranksum* in MATLAB). All tests were two-tailed.

To measure trends of latencies and activity amplitude across depths, we fitted a cubic function to the data of the form (*ax*^3^ + *bx*^2^ + *cx* + *d*), where *x* is the depth within SC measured using channel index ^15^. We chose higher order fits to capture non-monotonic trends across depths. *R*^2^ values were reported to assess the goodness-of-fit. A P-value for each cubic fit was obtained using a permutation test. For each permutation, the index of the SC depth was shuffled and a new shuffled-*R*^2^ value was obtained by fitting a new cubic function. 1000 permutations were done. The P-value was computed as the number of time shuffled-*R*^2^ values were superior or equal to the un-shuffled *R*^2^ value, divided by the number of permutations. A P-value inferior to 0.05 indicated a significant fit and, hence, that the data displayed a significant trend across depths.

## Acknowledgements

Funding for this research was provided by NIH grants R01 EY022854 and R01 EY024831.

## Author contributions

Design of study: CM, NJG; Experimental Data acquisition: CM, UKJ; Data analysis: CM; First draft of paper: CM; Writing and editing: CM, NJG; Final edits and proofing: CM, NJG, UKJ

## Materials & Correspondence

Corentin Massot

## Data and code availability

All data and code are available upon requests to Corentin Massot

---

**Supplementary Figure 1.**
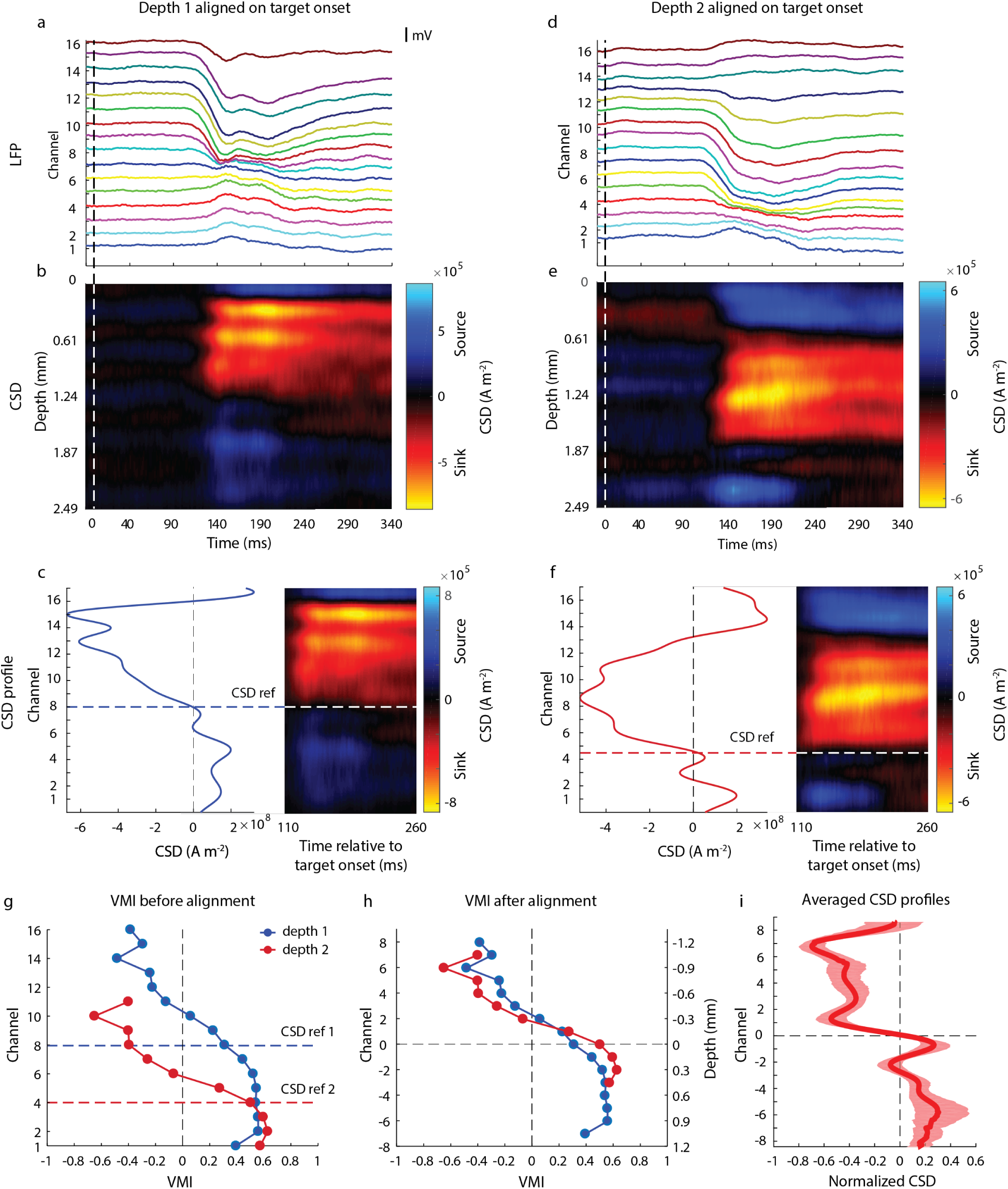
Alignment procedure based on LFP and CSD profiles. (a) Plots of trial-averaged LFP signals for one recording session with a 16-channel linear probe during a VG task; signals have been offset vertically to distinguish the activity on the different channels; channel 1 and 16 correspond to the most ventral and most dorsal contacts, respectively. Data are aligned on target onset. (b) Plot of CSD signal obtained from the LFP signals in (a), obtained with the iCSD method in the *csdplotter* toolbox (https://github.com/espenhgn/CSDplotter) ^78^. Negative and positive values correspond to current sinks and sources, respectively. (c) Another representation of the CSD profile (left panel). It shows the temporal average of the CSD values computed over the 150ms window following the sink onset (right panel). The horizontal line represents the depth of the crossing from negative to positive CSD values. This crossing is taken as the depth of reference for the alignment procedure. (d-f) Figures resulting from the same analysis as (a-c) for data recorded during the same session and for the same penetration but with the probe ∼1mm shallower in SC. (g) The VMI as a function of depth for the two examples presented in (a) (depth 1 in blue) and (d) (depth 2 in red); the horizontal dashed lines indicate the CSD reference channels. (h) The VMI traces of the two examples are replotted after alignment based on the CSD analysis; the channels’ index is reported on the left y axis. Channel 0 corresponds to the reference channel. The right y axis indicates the relative depth in mm. (i) The trace shows the CSD profile averaged across all sessions, not just the two examples from above, and for both VG and MG trials. The red translucid region surrounding the average CSD profile represents 95% confidence interval.

**Supplementary Figure 2.**
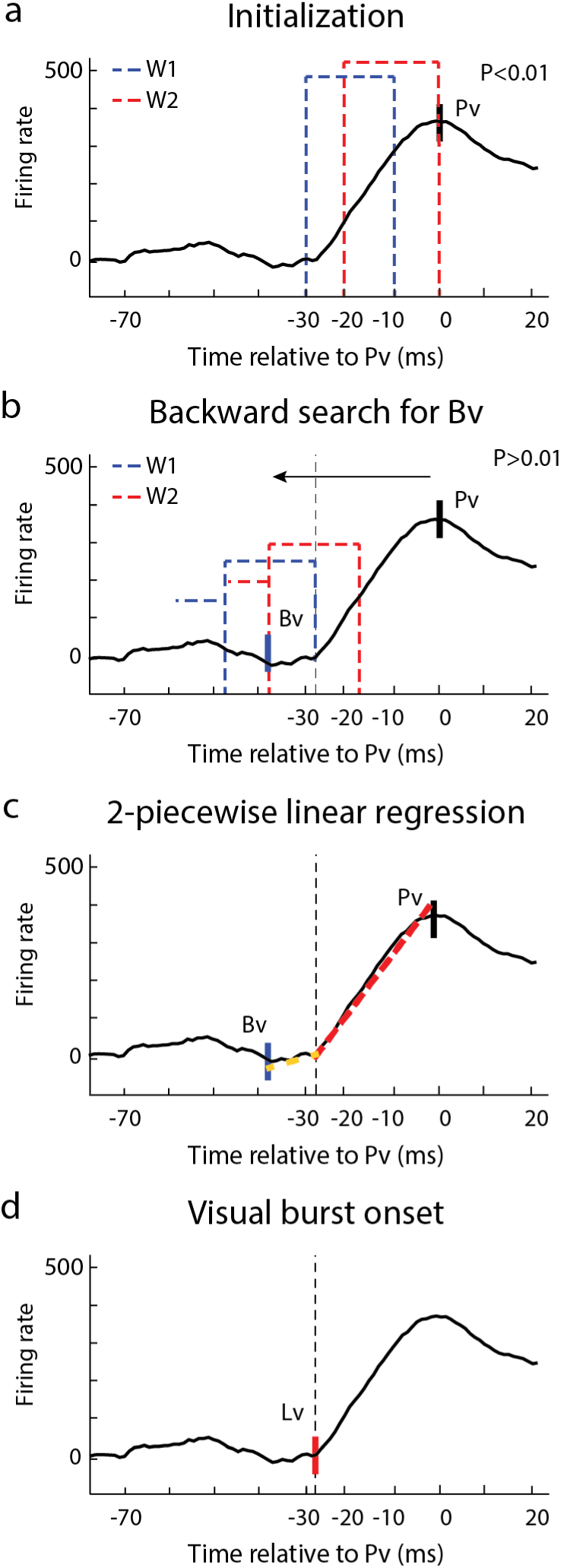
Visual latency detection. (a) Baseline-corrected average spike density waveform (solid black trace). *Pv* is the peak activity detected in the [-50 150]ms epoch after target onset. The x-axis represents time relative to Pv for display purposes. The two sliding windows *W1* (dashed red) and *W2* (dashed blue) are represented at their initial positions; a statistical difference between the distributions of activity in *W1* and *W2* is measured using a t-test; the test is significative at the initialization (P<0.01). (b) Both windows slide to earlier times in 1ms steps; the t-test is performed at each time step; *Bv* is the first time point (measured relative to the beginning of *W1*) when the *t*-test indicates a non-significant difference and stays not significant for the next 10 steps. (c) A two-piecewise linear regression is computed between *Bv* and *Pv.* (d) *Lv* is the time point that minimizes the residuals of the two-piecewise linear regression and represents the onset of the visual burst.

**Supplementary Figure 3.**
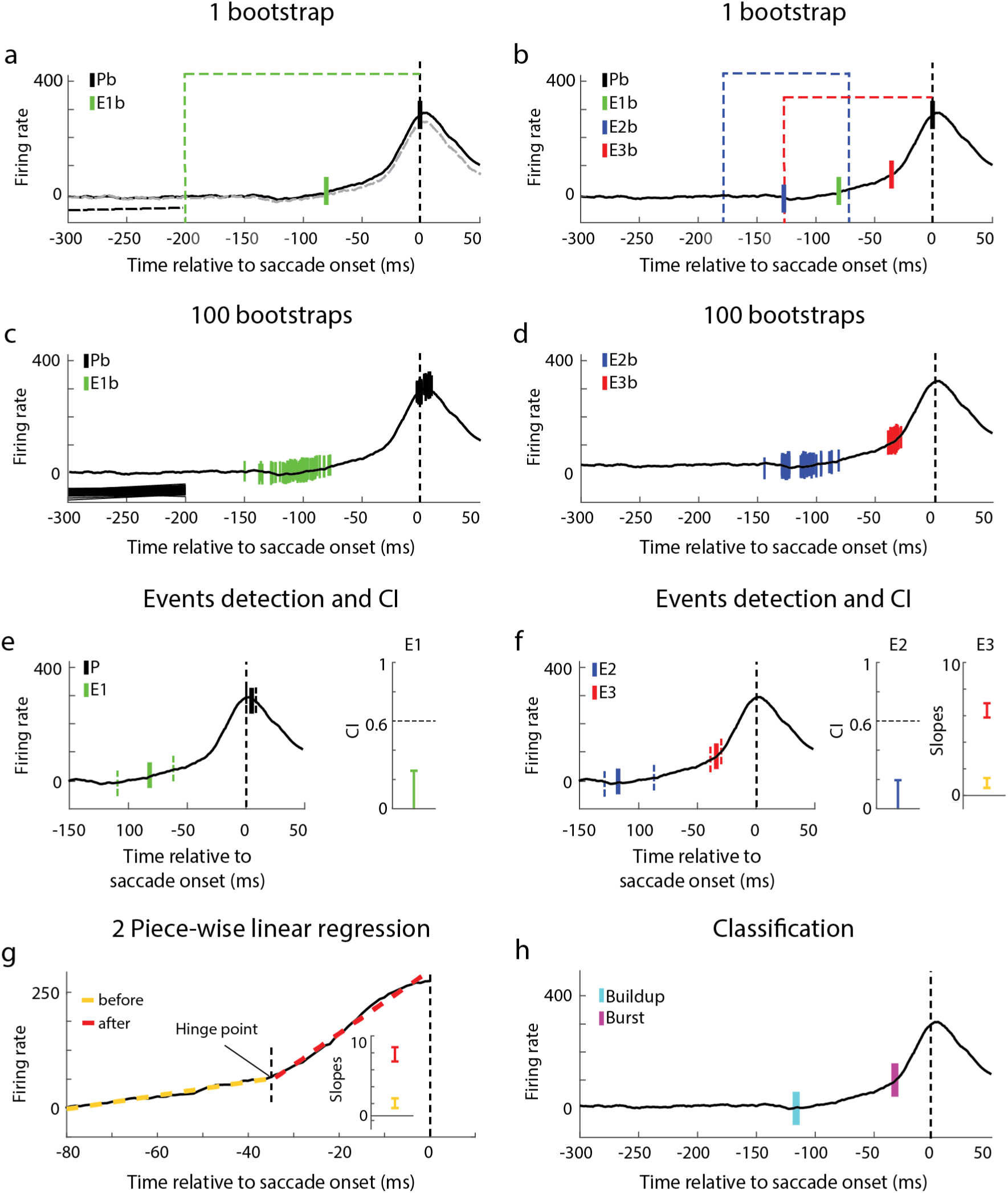
Events detection and classification of activity during the pre-saccadic epoch. (a) Average spike density waveform for one channel is shown for one bootstrap iteration (solid black trace). The near-horizontal, dashed black line is the linear trend estimated in [-300 −200]ms window. This trend was extrapolated to the remaining time points and subtracted from the trial-averaged activity to yield the dashed gray waveform. The time and amplitude at peak activity (*Pb*) and when the detrended activity becomes significantly different from baseline (*E1b*) for this bootstrap iteration are shown in black and green tick marks, respectively. The green dashed region delimits the search window of this event. (b) The original spike density waveform (black traces) now also overlays the ‘hinge’ points denoting *buildup* (*E2b*; blue tick mark) and *burst* (*E3b*; red tick mark) onsets for one bootstrap iteration. The blue and red dashed lines mark the search windows of the two events. (c,d) The spike density waveform is now shown with the distribution of each event for 100 bootstrap iterations. Subplots are separated for visualization. (e) Left: The mean estimates and confidence intervals (CIs) of events *P* (black) and *E1* (green) obtained from the 100 bootstrap iterations are shown respectively as solid and dashed tick marks. They are superimposed on the trial-averaged spike density function. Right: The normalized range of CI for *E1* is shown relative to an arbitrarily chosen threshold level indicated by the horizontal dashed line. (f) Left: Same format is used to shown the mean and CIs for events *E2* (blue) and *E3* (red). Middle: The normalized range of CI for *E2* is shown relative the same threshold level indicated by the horizontal dashed line. Right: The plot shows the means and CIs of the slopes of the regression fits before (yellow) and after (red) the hinge point of event *E3*. (g) A visualization of a hinge point detection from a two-piece linear regression analysis. The dashed lines indicated the best fit lines before (yellow) and after (red) the hinge point. (h) The final step of the analysis is to classify events *E2* and *E3* into *buildup* (cyan) and *burst* (purple) events (see *Materials and Methods* for criterion details).

